# Systematic Plasmid Engineering for Targeted Carotenoid Synthesis in Bacteria

**DOI:** 10.1101/2024.12.23.629938

**Authors:** Maiko Furubayashi

## Abstract

Carotenoids are natural pigments known for their vibrant colors and roles in various biological functions, such as antioxidant activity, photosynthesis, and radiation tolerance. The microbial production of carotenoids through heterologous expression of biosynthetic genes on engineered plasmid has facilitated the exploration of these functions. However, the complexity of creating carotenoid pathway plasmids has limited the number of available constructs, despite the vast diversity of carotenoids found in nature. This study aims to overcome these challenges by developing a platform for the systematic creation of plasmids that efficiently produce specific carotenoids in bacteria. A novel strategy, called Slot Assignment Cloning (SA-Clo), which applies the modular cloning (MoClo) method, was introduced. Using this strategy, plasmids for 17 carotenoids were constructed, including lycopene, β-carotene, zeaxanthin, canthaxanthin, astaxanthin, phytoene, δ-carotene, ε-carotene, α-carotene, violaxanthin, isorenieratene, several C_30_ carotenoids, and non-natural C_50_-carotenoids. These plasmids enable the targeted and specific production of carotenoids, streamlining future studies on lesser-known carotenoids and their applications.

## Introduction

Carotenoids are natural pigments known for their vibrant colors, with over 850 kinds identified so far.^1,2^ They are known for photosynthesis and photoprotection in plants, or for antioxidant effects and vitamin A provision for animals.^3^ Accumulating research in the past decades indicates that the scope of carotenoid functions extends significantly beyond these traditional roles—for instance, some carotenoids can act as an antenna for a protein photo-switch^4^ (canthaxanthin, **Fig. 1**) or for a light-driven proton pump^5^ (salinixanthin), or even play a role in radiation tolerance for the most radiation-resistant microbe discovered^6,7^ (deinoxanthin). The carotenoids also can be a precursor of “apocarotenoids”, a group of compounds derived from the oxidative cleavage of carotenoids, which play crucial roles in various biological processes, including signaling, pigmentation, and stress response.^8^ In addition, different families of soluble proteins that binds to a specific kind of carotenoids are discovered from different organisms, including lobsters,^9^ cyanobacteria^4,10,11^, human^12^ and silk moth^13,14^; their functions are now under investigation with great interest. These discoveries further ensure the view that carotenoids are not just a passive molecule located in the membrane, but they are rather involved in diverse form and functions.

**Figure 1.**
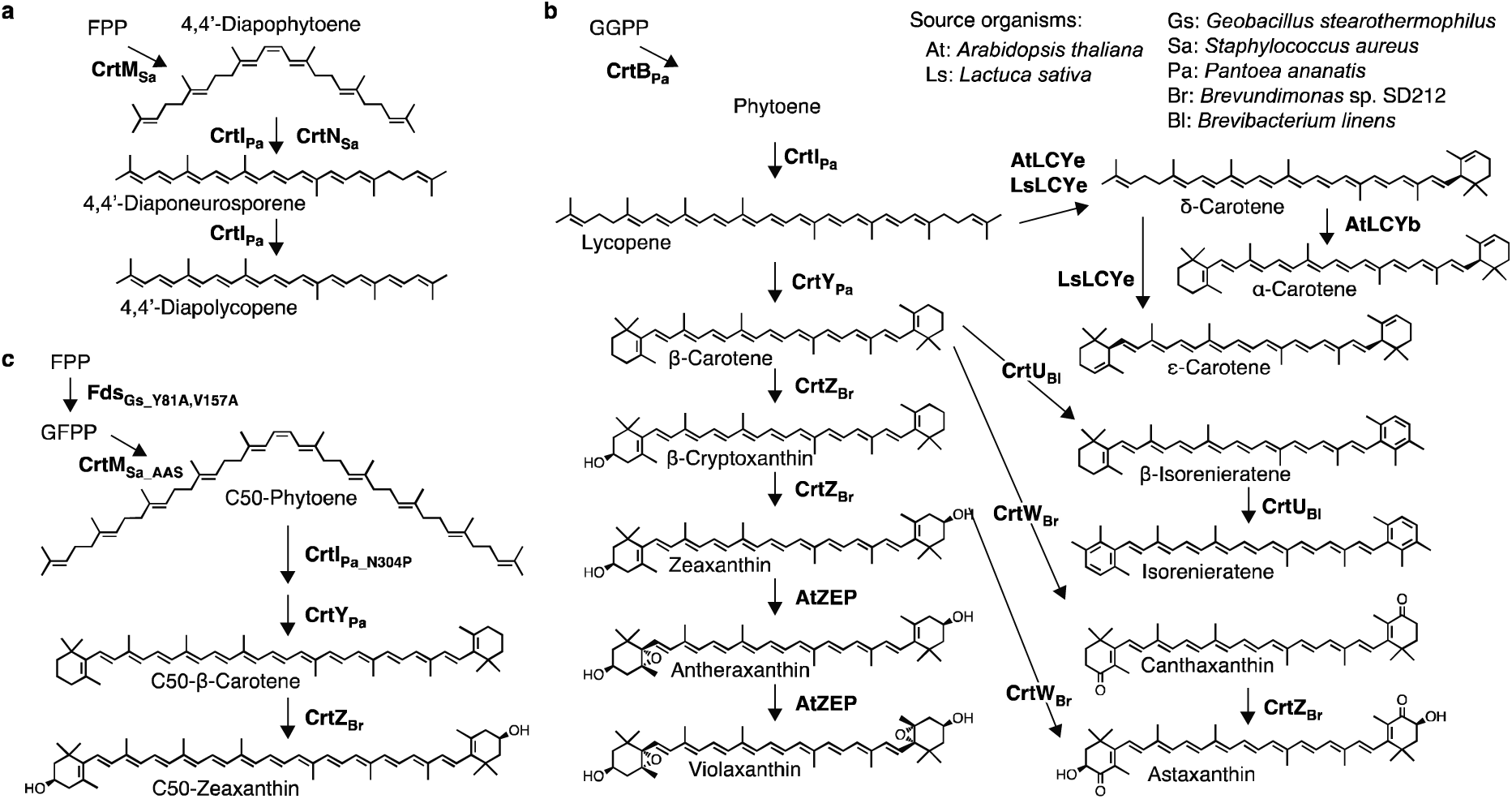
Carotenoid biosynthetic pathways reconstructed in this study. The source organism for each enzyme is indicated by a suffix and subscript (for bacteria) or by a prefix (for plants), corresponding to the organisms listed in the top-right corner. (a) Carbon-30 (C_30_) carotenoid pathway, showing the biosynthesis of C_30_ diapocarotenoids found in some bacteria. (b) C_40_ carotenoid pathway, showing the biosynthesis of common carotenoids such as lycopene and β-carotene. (c) Non-natural C_50_ carotenoid pathway, which was reported in a previous study.^27^ Abbreviations: FPP, farnesyl diphosphate; GGPP, geranylgeranyl diphosphate; GFPP, geranylfarnesyl diphosphate; CrtM_Sa_AAS_, CrtM_Sa_ with F26A/W38A/F233S amino acid substitutions^27^; CrtI_Pa_N304P_, CrtI_Pa_ with N304P amino acid substitution.^27^

Using microorganisms like *Escherichia coli* to reconstruct/refactor these carotenoid-utilizing biological functions would be highly effective for the understanding or application of such functions. Examples of early studies include the reconstruction microbial rhodopsin in *E. coli* using a retinal (β-carotene cleavage product) pathway,^15–17^ or the reconstruction of photo-switching orange carotenoid protein (OCP) function in *E. coli* by the introduction of echinenone/canthaxanthin pathways.^18–20^ Creating additional line-up of heterologous carotenoids pathways could facilitate the reconstruction of many more carotenoid-utilizing biological functions.

Since the discovery of the carotenoid biosynthetic genes in soil bacteria,^21^ many plasmids for heterologous carotenoid production have been developed,^22,23^ and widely distributed via plasmid repositories such as addgene or RIKEN BioResorce Research Center (BRC), which hugely contributed to the progress of carotenoid and pathway engineering research. However, to date, the capacity to produce carotenoids as a sole product using a single plasmid in *E. coli* has been mostly confined to major carotenoids, such as lycopene, β-carotene or astaxanthin, majority of them found in plants. The heterologous production of relatively minor carotenoids is still limited, making it difficult to reconstruct the carotenoid-utilizing protein function that uses such carotenoids. Creating a more diverse array of carotenoid production plasmids enables their use in various reconstitution studies, enriching our understanding of carotenoid-using biological function and aiding in applied research. The requirements for such carotenoid plasmids are (1) to preferably all the gene is expressed by a single plasmid (since the carotenoid-binding-protein-of-interest would be able to be expressed by other vectors), and (2) to produce the target carotenoid as a sole product, with minimal byproduct or intermediates.

The rare carotenoids are especially difficult to produce in a targeted manner, since they often require multiple genes from different source organisms to reconstruct an efficient pathway, thereby creating the need to optimize the expression levels. For example, even the relatively simple carotenoid astaxanthin, which requires six genes to produce in *E. coli* (**Fig. 1**), has often been used as a model case to optimize the expression, since the simple co-expression of 6 genes often would not result in a sole product in *E. coli*.^24^ In addition, there have been many studies regarding optimizing multi-gene expression,^25^ but those strategies are often used for the target with deep interest, and requires significant effort for each optimization. For the construction of plasmids for various carotenoid production, a more quick-and-easy method to just obtain the sole product by *E. coli* would be useful.

This study aims to establish a platform for systematically creating plasmids for carotenoid production and used them to construct a series of plasmids that exclusively produce the target carotenoids. Initially, plasmids for six major carotenoids (lycopene, β-carotene, zeaxanthin, canthaxanthin, astaxanthin), important “hub” carotenoids, were constructed using step-by-step cloning. During this process, the necessity to modify gene types and order for optimal expression became apparent. For this purpose, the modular cloning (MoClo) method^26^ was applied for the creation of other carotenoid pathways. This system, named Slot Allocation Cloning (SA-Clo), facilitates rapid construction of operons with various constructs, enabling the specific production of various carotenoids including: phytoene, δ-carotene, ε-carotene, isorenieratene, α-carotene, 4,4’-diapophytoene, 4,4’-diaponeurosporene, 4,4’-diapolycopene and violaxanthin as well as non-natural carotenoids^27,28^ such as C_50_-phytoene, C_50_-β-carotene and C_50_-zeaxanthin. SA-Clo provides the capability to create plasmids for a variety of minor and rare carotenoids, which could significantly advance the study of diverse carotenoid-utilizing biological systems.

## Results and Discussion

### 1. Plasmid construction for the production of lycopene, **β**-carotene, zeaxanthin, canthaxanthin, and astaxanthin

To build a platform for various carotenoid operons, I first aimed to design and build plasmid constructs for five key carotenoids i.e. lycopene, β-carotene, zeaxanthin, canthaxanthin, and astaxanthin. The purpose for these plasmids was not to maximize carotenoid production but to achive a sufficient production levels (easy extraction, colony-color visibility and low toxicity, which is approx. 1-10 mg/L) while ensuring the biosynthesis of a single carotenoid product to minimize the formation of intermediates.

Given that simple co-expression of biosynthetic genes can often lead to an accumulation of intermediate compounds,^24^ careful control over expression levels is crucial for targeted production. From my previous studies^28–30,27^ and common sense, it has been observed that high enzyme activity (i.e. expression level and kinetics) in the later stages of the pathway prevents the accumulation of intermediates, thus facilitating the production of the intended carotenoid. Based on this knowledge, I developed guidelines to design these five carotenoid plasmids, which are elaborated in **Supplementary Note 1**. In short, the genes were arranged into an operon format, initiated by a single promoter. The promoter was immediately followed by the GGPP synthase gene (*ggps\**, see **Supplementary Note 1**) and phytoene synthase gene (*crtB*). The downstream enzymes genes (such as *crtI* and *crtY*) were arranged in reverse order of their action within the pathway, with genes for enzymes acting later in the pathway positioned closer to the promoter.

The above strategies were used to design and construct the plasmids for lycopene, β-carotene, zeaxanthin, canthaxanthin, and astaxanthin (**Figure 2a**). The plasmids were introduced into *E. coli*, cultured for two days and the extracted carotenoids were subjected to liquid chromatography (LC) analysis. For each construct, it was confirmed that intended carotenoids were the major product, with negligible intermediates (**Figure 2b**). Furthermore, since these operons are all under a single promoter, I investigated whether altering the promoter could change the production levels. Instead of P_J23115_ (https://parts.igem.org/Part:BBa_J23115), the promoters with different strengths (P_J23101_, P_J23107_, P_J23117_) were used and the production levels were analyzed. As expected, the carotenoid production levels resulted in the order of promoter strength (**Figure 2c**).

**Figure 2.**
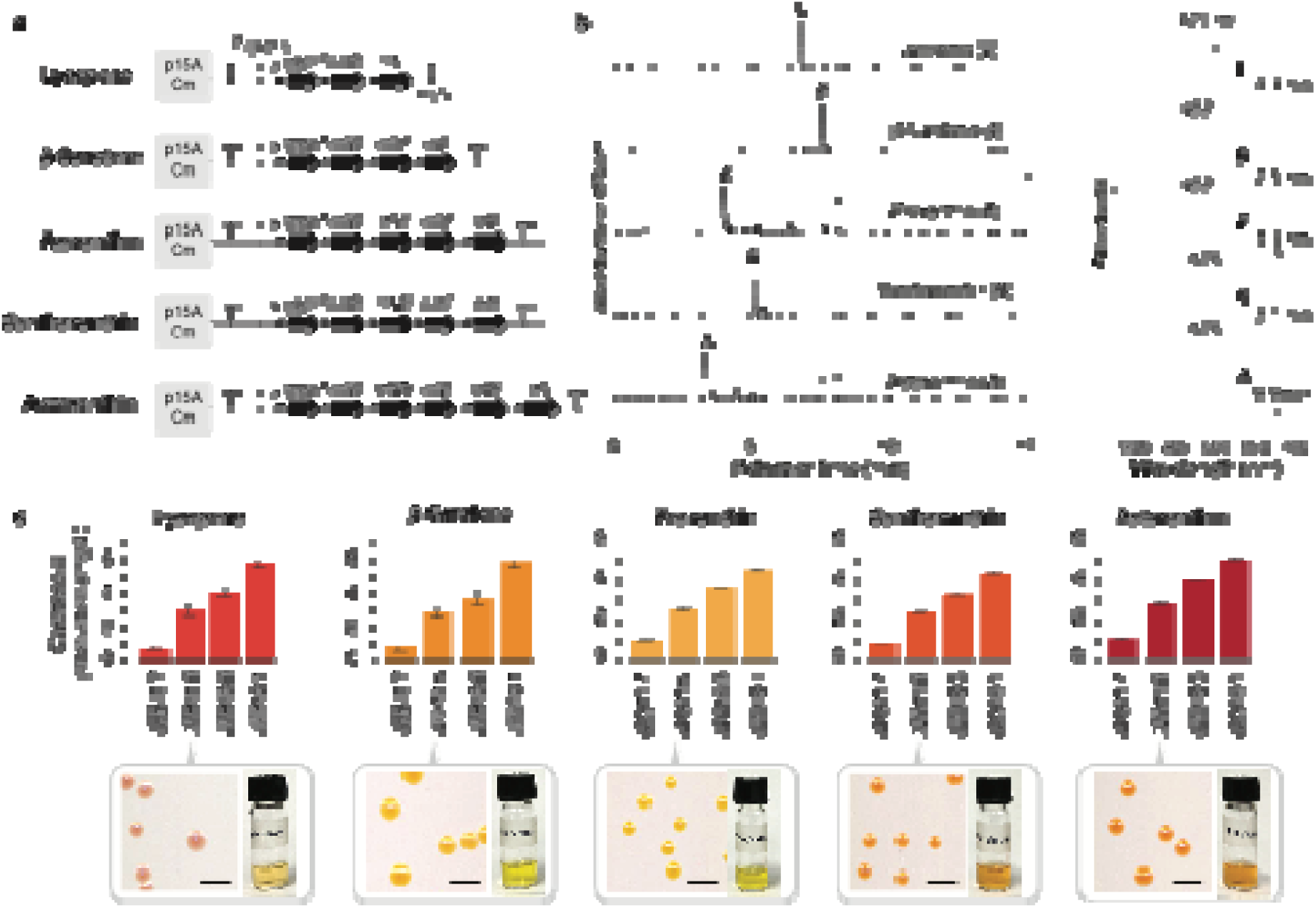
Plasmid construction for “major” carotenoid pathways. (**a**) Plasmid constructs for each carotenoid pathway. (**b**) LC-PDA chromatogram of carotenoid extracts from E. coli harboring the indicated plasmids from panel **a.** The absorbance spectrum of each peak is shown on the right. (**c**) Carotenoid production titer (in mg/L) by E. coli harboring plasmids indicated in panel **a**, but with different promoters. See Supplementary Table 1 for the plasmids used in this figure. The bars indicate the average from 4 samples, and the error bars indicate the standard deviation. The *E. coli* colony (scale bar 0.5 mm; plated onto nitrocellulose membrane to provide white background) and carotenoid extracts (in 1 mL acetone) shown on the bottom are from the cells harboring constructs using the P_J23115_ promoter. Abbreviations: A, astaxanthin; C, canthaxanthin; L, lycopene; Z, zeaxanthin; β, β-carotene.

### 2. Development and Architecture of the System for “Slot Assignment Cloning”

While the five major carotenoid plasmids were constructed successfully, to expand the plasmid line-up, there was a need for a more efficient method than the cumbersome task of individually rearranging and inserting genes. To address this, I adopted the Modular Cloning (MoClo) methodology.^26^ MoClo utilizes the cloning technique using a type IIS restriction enzyme (such as GoldenGate Cloning^31^) for creating DNA constructs in a monosistronic “transcription units” in a modular and streamlined manner, and have been demonstrated in a diverse organisms, including bacteria, fungi, plants and mammalian cells.^32^ The original MoClo involves preparing a wide range of gene regulatory parts, such as promoters and terminators, in Level-0 plasmids, and assembling them with target genes to create Level-1 “transcription units”. These units are then combined to construct Level-2 plasmids. Though effective for studies requiring monocistronic expression, this method is not ideal for carotenoid plasmid construction for bacteria, where the use of numerous promoters and terminators is unnecessary and unwieldy.

The new system in this study named “Slot Assembly Cloning (SA-Clo)” was built upon the foundational principles of MoClo, tailoring it specifically for the synthesis of the carotenoid operon. SA-Clo retains the core cloning mechanism of MoClo, but it diverges in its design philosophy. Here, SA-Clo introduces a novel Level-1 vector design featuring ‘arrays’ with empty ’slots’ (**Figure 3**). These slots allow for dynamic rearrangement of gene sequences, catering to the specific needs of carotenoid construct assembly. This approach facilitates the seamless integration into Level-2 vectors, significantly enhancing the flexibility and efficiency of the carotenoid construct assembly process, compared to the conventional method in which one needs to design and order oligo primers.

**Figure 3.**
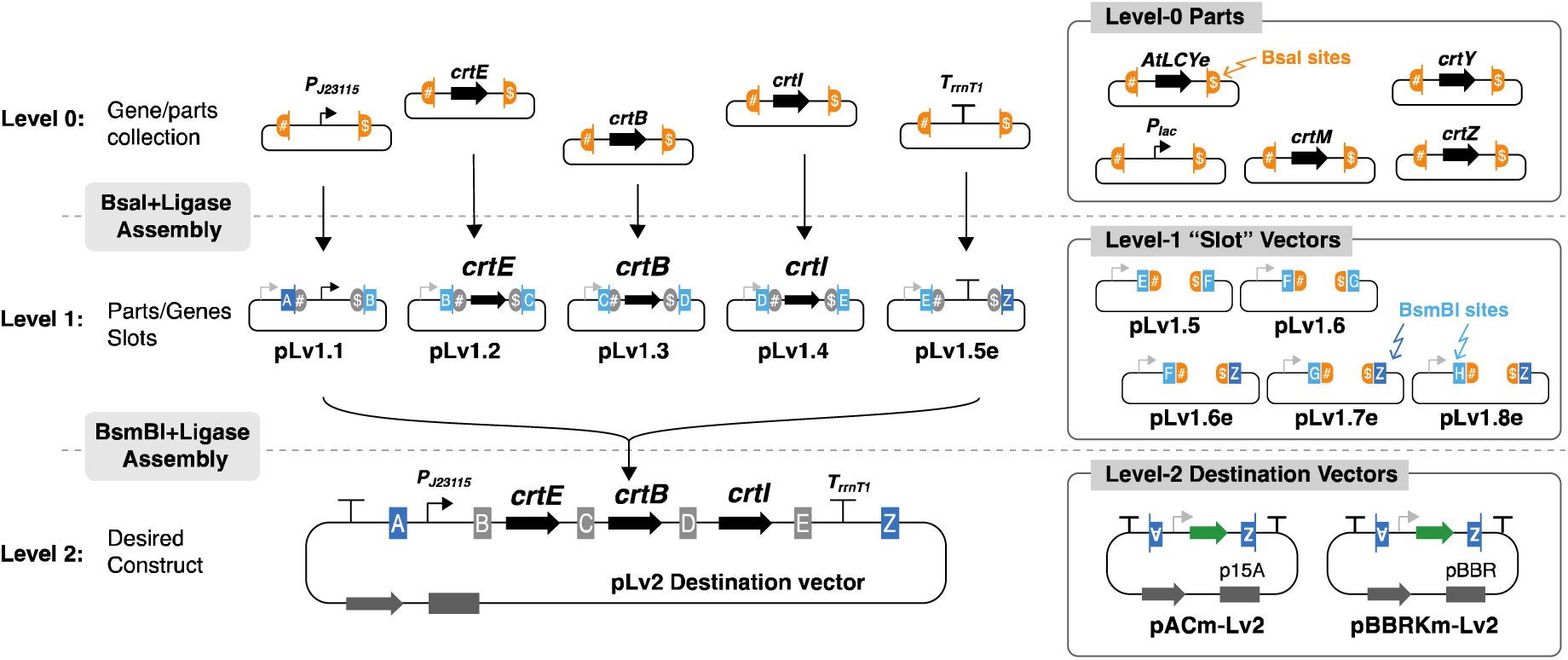
Overview of the carotenoid plasmid construction using Slot Assignment Cloning (SA-Clo). Level-0 plasmids harbor the enzyme genes and genetic parts such as promoters or terminators. The Level-0 genes/parts are individually inserted into the Level-1 vectors which work as “slots” via BsaI GoldenGate Assembly. The Level-1 plasmids are then assembled using BsmBI GoldenGate Assembly into a Level-2 destination vector.

Figure 3 provides an integrated schema of the SA-Clo framework. The initial step involved curating a comprehensive gene library for carotenoid-related genes and assorted genetic components within the Level-0 vector (Figure 4a). The Level-0 vector was equipped with a BsaI-site to facilitate seamless integration and included a GFP expression cassette useful for identifying the vector background. A diverse array of carotenoid enzyme genes and genetic elements, sourced from various organisms, were then introduced into the Level-0 vectors.

**Figure 4.**
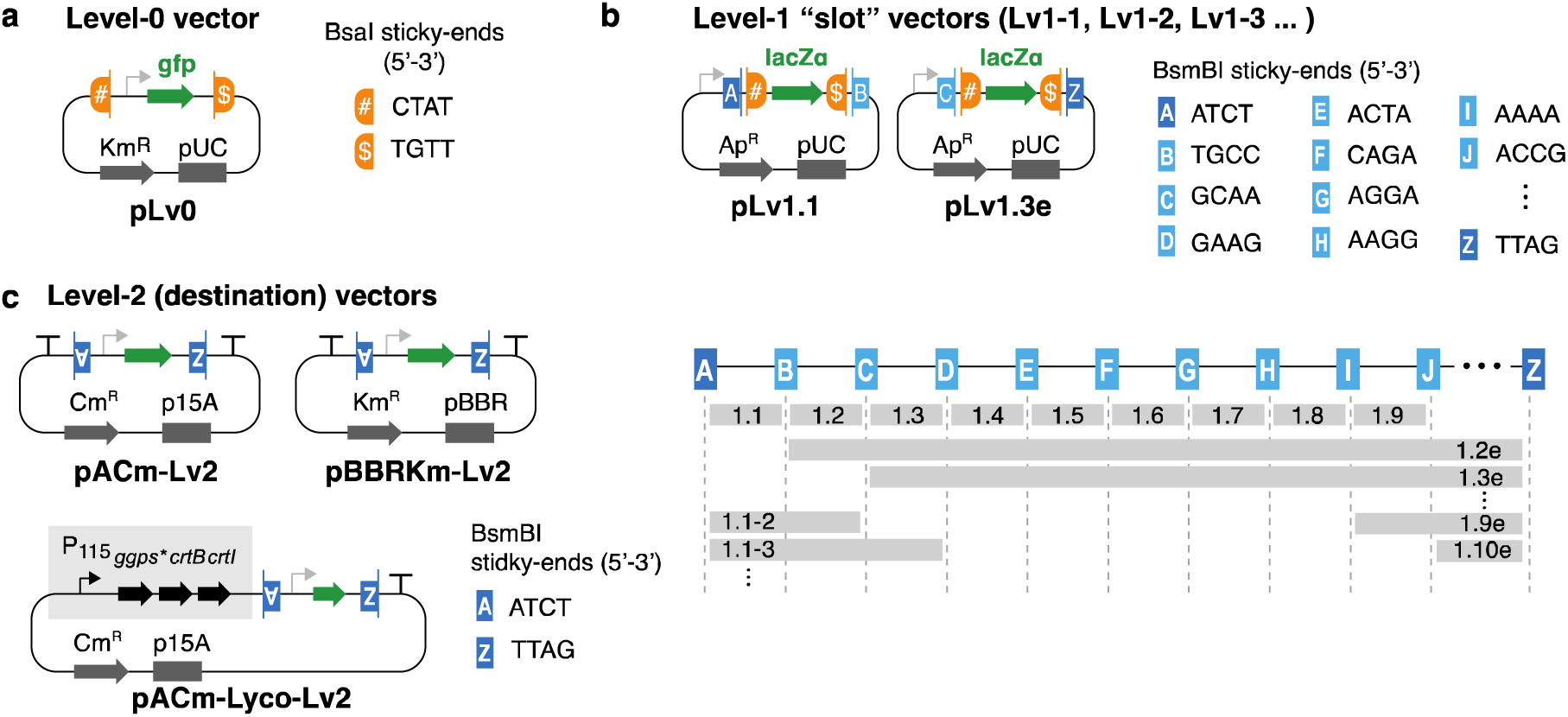
Vectors for Slot Assignment Cloning (SA-Clo). (**a**) Level-0 vector. Carotenoid enzyme genes or genetic parts (promoters or terminators) are held in this vector (see **Figure 3**). (**b**) Representative examples of Level-1 vectors. These vectors act as “slots” with pre-determined order for the genes/parts to be inserted. On the right, the BsmBI sticky-ends are indicated. On the bottom, the location of each “slots” are indicated. (**c**) Representative examples of Level-2 destination vectors. They include the front- and last-end of BsmBI sticky-ends (indicated by letter A and Z).

When the genes were ordered to a DNA-synthesis company, we chose the pUC/Km vector, and sequences were flanked with BsaI sites specific to Level-0 (see pLv0 sequence in **Supplementary Table 2**). For the RBS, I initially designed strong RBSs for each gene by using RBS calculator^33^ or UTR-designer.^34^ However, since my go-to RBS sequence (AGGAGGATTACAAA) appeared to be effective for many genes, I eventually used this sequence as a default. This sequence seemed to work adequately for most of the constructs (see Discussion). The assemblage of the Level-0 plasmids in this study is listed in **Supplementary Table 3**.

Next, these Level-0 components were integrated into the Level-1 vectors (Figure 4b). Each Level-1 vector was configured with designated sites for BsaI-mediated insertion of Level-0 parts, complemented by BsmBI sites to facilitate the transition to Level-2 assembly (Figure 3 **and 4c**). Between the BsaI sites are *lacZ* expression cassette, which would be useful for identifying vector backgrounds using blue-white screening. The order of the Level-1 vector was pre-determined, with slots for positions such as Level-1.1, Level-1.2, Level-1.3, etc. (Figure 4b). Using the NEBridge GetSet® tool (New England Biolabs), I created 11 different sets of unique overhangs that is predicted to result in high fidelity assembly, resulting in the capacity to assemble up to ten genes (Figure 4b). To provide flexibility in the number of parts to be assembled, “end-linkers” were incorporated into every slot (e.g., Lv1.3e, where “e” indicates an end-linker). This is based on the concept of the original MoClo’s end-linker.^26^

For example, three genes/parts could be assembled using Level-1.1, 1.2 and 1.3e plasmids, while five genes/parts could be assembled using Level-1.1, 1.2, 1.3, 1.4 and 1.5e plasmids. Level-1.1 is typically designated as a promoter slot, while the end-linkers often serve as terminators slot (“Term” in **Supplementary Table 3**). Accordingly, a variety of Level-1 plasmids with multiple promoters in Level-1.1 and terminators in Level-x.1e (where x is a numeral) were prepared.

Finally, the prepared Level-1 vectors were assembled into the Level-2 destination vector using BsmBI Golden Gate assembly (Figure 4c). For Level 2 destination vectors, not only empty vectors (pACm-Lv2, pAKm-Lv2) but also a range of carotenoid operon plasmids (pACm-lyco-Lv2, pACm-beta-Lv2, pACm-cantha-Lv2, pACm-zea-Lv2 and pACm-asta-Lv2) were prepared. Since many carotenoids uses these “hub” carotenoids as the pathway intermediate, these carotenoid Level-2 vectors could be beneficial in constructing such plasmids .

### 3. Testing the efficiency of type IIS GoldenGate reaction of SA-Clo System

Having the vectors prepared, first, the efficiency of the Level-1 assembly (BsaI GoldenGate assembly) was examined. As a test case, Level-0 parts (P_J23105_ promoter) were inserted into Level-1.1 vector (pLv1.1). The two plasmids (pLv0-P_J23105_ and pLv1.1) were mixed with NEB Goldengate BsaI kit in total 10 µL reaction (see Methods). The reaction was either performed with 37°C 5 min + 60°C 5min (recommended for cloning in the manufacture instruction) or 37°C 60 min + 60°C 5 min (recommended for library construction). The reaction mixture was used for transformation of E. coli competent cells, and a portion of the cells were plated on LB-agar supplemented with X-gal, IPTG and antibiotics blue-white screening.

As a result, for both the 37°C 60-min and 37°C 5-min reaction sample, no vector background (blue-colored colony by X-gal) was observed (**Supplementary** Fig. 1). The 5-min reaction resulted in 25 cfu/transformation, while the 60-min reaction resulted in 3000 cfu/transformation. Colony PCR confirmed that all colonies contained the intended insert (**Supplementary** Fig. 1c). Additionally, due to the high efficiency of the BsaI assembly, the process was streamlined by culturing transformants (SOC culture) directly in a fresh LB liquid media followed by overnight culture and plasmid miniprep, substantially reducing the time involved (detailed in the Methods section). The 5-min reaction transformants resulted in 2.5 µg/50 µL miniprep solution from 2-mL culture, while the 60-min transformants resulted in 5 µg/50 µL of plasmid DNA from 2-mL culture.

Next, as a test case for Level-2 assembly, I chose canthaxanthin as a target since the color is easy to visualize. To this end, five distinct Level-1 plasmids were created: pL1.1-Para, pLv1.2-ggps*, pLv1.3-crtB, pLv1.4-crtW, pLv1.5-crtY, pLv1.6-crtI, pLv1.7e-Term (**Supplementary** Fig. 2). For the assembly process, I used the above seven Level-1 plasmids and pACm-Lv2 (Level-2) vector, and mixed with NEB BsmBI GoldenGate assembly kit for 45°C 1min, 16°C 1 min for 40 cycles, followed by 60°C 5min, following the manufacture’s instruction. The results show that there were 0% vector background (GFP colony), and the 40 of the 42 colony showed an orange color, indicating a high success rate (**Supplementary** Fig. 2). Four orange colonies were picked, miniprepped, and digested by restriction enzymes for confirmation, verifying the results. Notably, this success rate was achieved despite non-optimized conditions: the plasmid DNAs used were 0.5 µL (ranging from 50-150 ng), the molecular ratios were not carefully calculated as recommended in the GoldenGate kit, and the competent cells used were only 10 µL instead of the frequently used 20-50 µL for transformation. This demonstrates the ease and robustness of this system.

### 4. Construction of several carotenoid operon plasmids by SA-Clo and characterization

Using the above established SA-Clo system, I have embarked on the construction of several carotenoid operons, focusing on the synthesis of carotenoid beyond the initial set (lycopene, β-carotene, zeaxanthin, canthaxanthin and astaxanthin) that had already built in the above sections in Figure 2. The goal here was to construct plasmids for the production of various carotenoids by using the genes are well known in other literature. These include 12 carotenoids: 5 carotenoids well known in plants (δ-carotene, ε-carotene, α-carotene, violaxanthin and phytoene), 4 microbial carotenoids (isorenieratene, 4,4’-diapophytoene, 4,4’-diaponeurosporene, 4,4’-diapolycopene), and 3 non-natural long-chain C_50_-carotenoids^27^ (C_50_-phytoene, C_50_-β-carotene, C_50_-zeaxanthin, all created in the laboratory by directed evolution).

The 12 plasmid constructs were designed for each carotenoid and assigned the Level-1 and Level-2 plasmids to be constructed (**Supplementary Table 4**, top 12 rows). Since it is difficult to rationally design the optimal expression balance, I simply decided the gene order and parts selection by guesswork (and by following some guidelines in **Supplementary Note 1 (2)**) to be a starting point for further optimization. The plasmid was assembled within 1 week (2 days for Level-1 assembly and 3 days for Level-2 assembly and sequence confirmation). *E. coli* strains harboring the constructed plasmids were cultured in liquid media and their carotenoid production was characterized.

Among the 12 plasmids, 8 constructs predominantly yielded the target compounds (constructs shown in Figure 5). The LC-PDA chromatogram of the *E. coli* acetone extracts all resulted in single peak for the target carotenoids. The color of the colonies and acetone extracts resulted in vibrant yellow or red (except for the colorless carotenoids, phytoene and its analogues). The results demonstrate the power of SA-Clo: the process was very quick and easy compared to the conventional carotenoid plasmid construction, where one designs the construct from scratch and orders DNA fragments/oligo, followed by performing different DNA assembly process each time. Since all the assembly step is streamlined into only Level-0, Level-1, or Level-2 assembly, many different carotenoid plasmids could be constructed in parallel without any complication or difficulty during the process.

**Figure 5.**
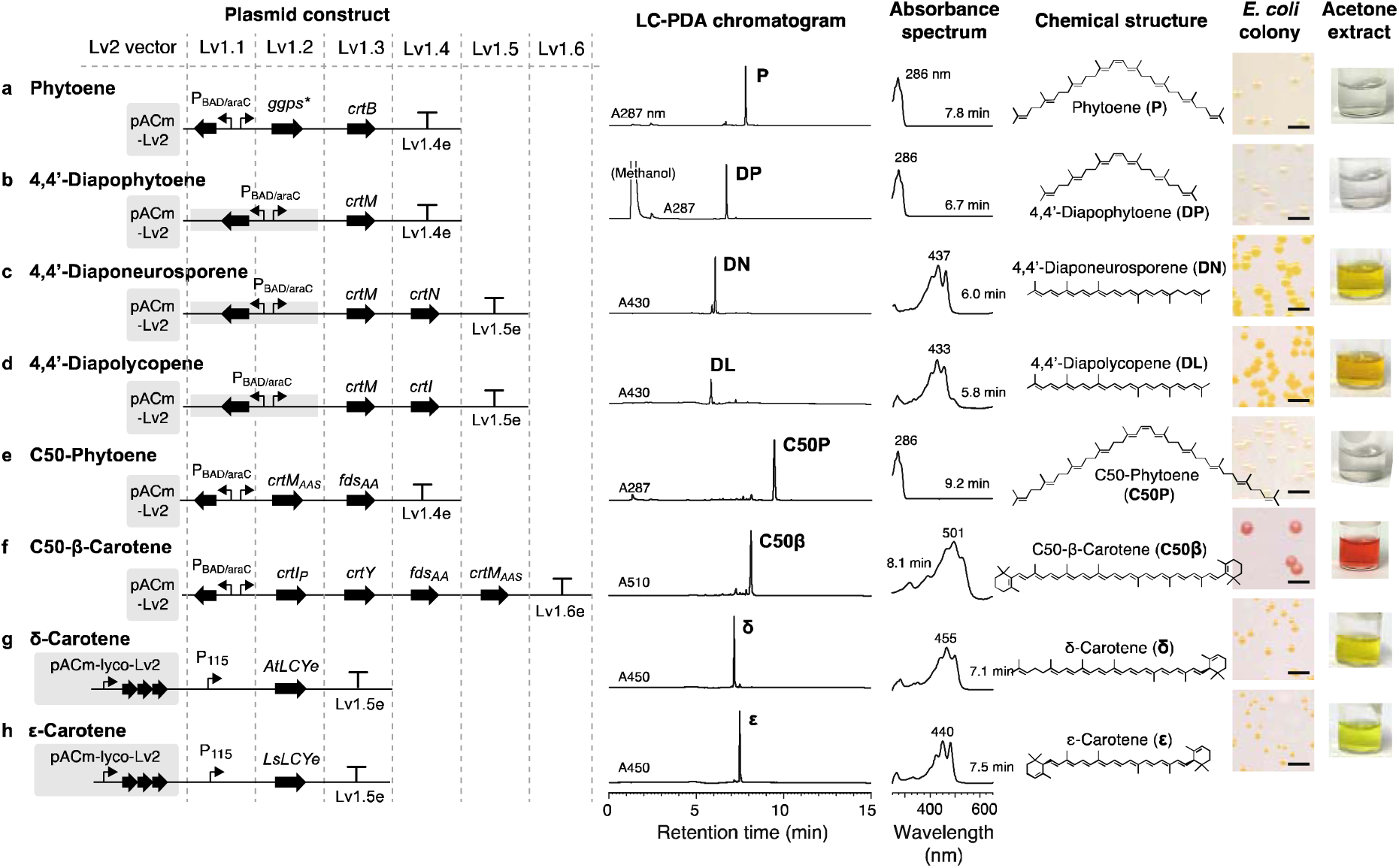
Various carotenoid plasmids constructed by SA-Clo that resulted in selective production of the target carotenoid. From left to right, plasmid constructs, LC-PDA chromatogram of carotenoids extracted by *E. coli* (monitoring wavelength in nm indicated after “A”), absorbance spectrum of each peak, the chemical structure of the target carotenoid, *E. coli* colony (scale bar 0.5 mm, plated onto a nitrocellulose membrane to provide white background) and photo of the 1 mL-acetone extract are shown. (**a**) Phytoene (P); (**b**) 4,4’-diapophytoene (DP); (**c**) 4,4’-diaponeurosporene (DN); (**d**) 4,4’-diapolycopene (DL); (**e**) C_50_-phytoene (C50P); (**f**) C_50_-β-carotene (C50β); (**g**) δ-carotene (δ); (**h**) ε-carotene (ε).

The plasmids constructed for the remaining four carotenoids (isorenieratene, violaxanthin, α-carotene and C_50_-zeaxanthin) resulted in mixed products, as indicated in Figure 6a ***(i)*, 6b *(i)*, 6c *(i)*** and **6d *(i)***, respectively. These candidates were subjected to the next round of design and construction.

**Figure 6.**
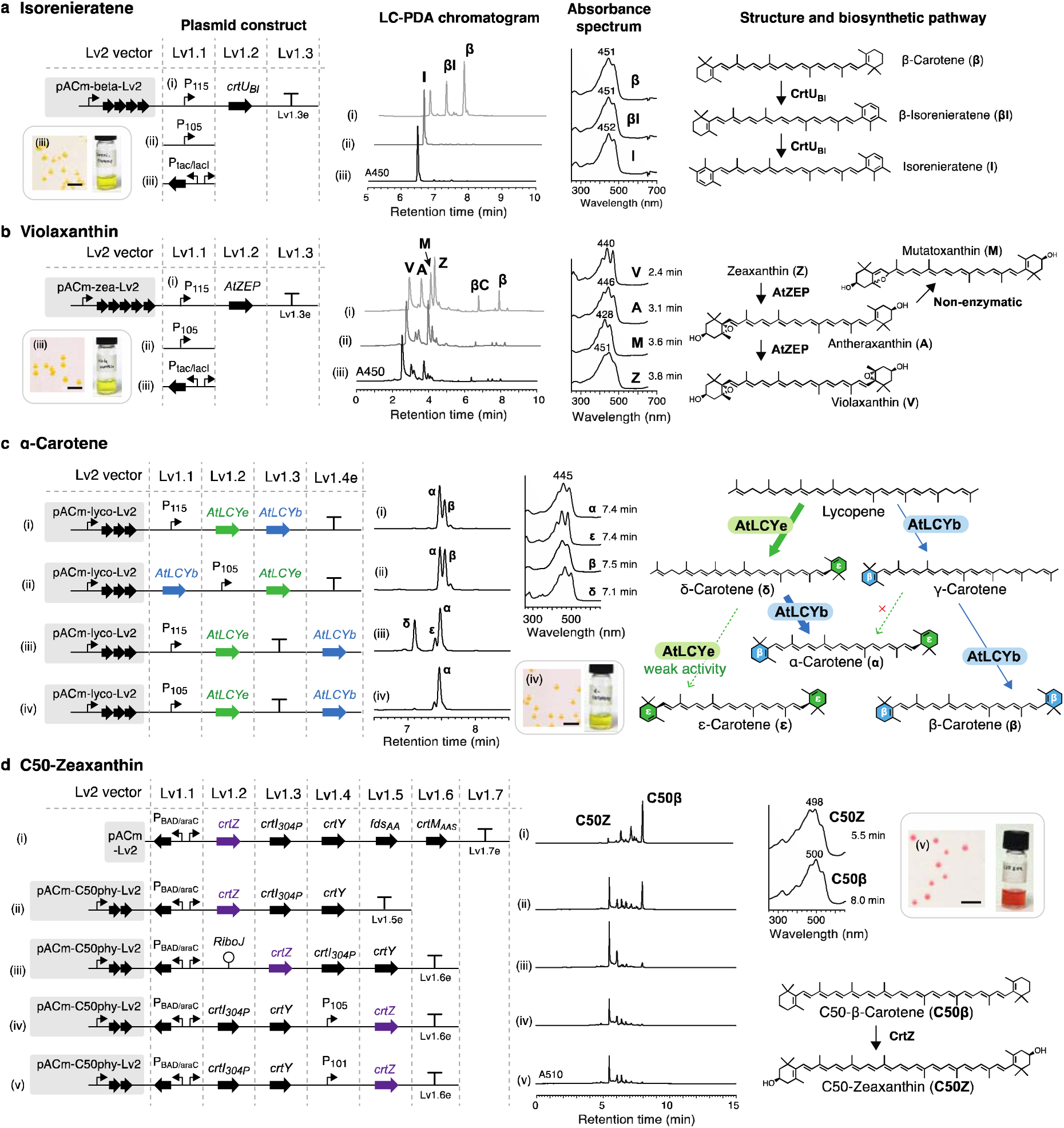
Optimization of carotenoid plasmid construction using SA-Clo. From left to right: the plasmid constructs, LC-PDA chromatogram of E. coli carotenoid extracts, absorbance spectrum of each peak, and the structure and biosynthetic pathways are shown. The colony color and acetone extract are shown as insets. (**a**) Plasmid construction for the targeted production of isorenieratene. Three constructs with different promoters in Level-1.1 are indicated by i, ii and iii. They correspond to the three LC-PDA chromatograms at 450 nm. (**b**) Plasmid construction for the targeted production of violaxanthin. Three constructs with different promoters in Level-1.1 are indicated by i, ii and iii. They correspond to the three LC-PDA chromatograms at 450 nm. Mutatoxanthin are generated by a non-enzymatic reaction.^29^ Abbreviations: βCr, β-cryptoxanthin, β, β-carotene. (**c**) Plasmid construction for the targeted production of α-carotene. Four different constructs (i, ii, iii, iv) are indicated. On the right, the pathway towards α-carotene is indicated in thick arrow. α-Carotene is biosynthesized via δ-carotene, but not γ-carotene. (**d**) Plasmid construction for the targeted production of C_50_-zeaxanthin. Five constructs are build indicated as i, ii, iii, iv and v. aiming to optimize the crtZ gene expression (see main text). The LC-PDA chromatograms at 510 nm are shown.

For isorenieratene, the initially designed construct (Figure 6a ***(i)***) resulted in not just the intended isorenieratene, but also the accumulation of the pathway intermediates (β-carotene and β-isorenieratene). This indicated that the initial promoter choice (P_J23115_) was not suitable and that a stronger promoter was required. By creating a new construct with a stronger promoter (P_J23105_ and P_tac/lacI_), more efficient accumulation of isorenieratene was achieved (Figure 6a *(**ii)*** and ***(iii)***).

Similarly, the initial violaxanthin construct (Figure 6b ***(i)***) resulted in a mixture of 6 carotenoids in addition to violaxanthin. The accumulation of pathway intermediate zeaxanthin and antheraxanthin indicated that the promoter strength (expression level) of AtZEP was insufficient for this case as well. Among the two additional constructs designed with stronger promoters (P_J23105_ and P_tac/lacI_), the one with P_tac/lacI_ promoter resulted in relatively high accumulation of violaxanthin (Figure 6b ***(iii)***). Mutatoxanthin was accumulated which may derived by a non-enzymatic reaction.^29^

For the plasmid designed for α-carotene, the initial construct resulted in a mixture of α-carotene and b-carotene (Figure 6c ***(i)***). It has been known that the heterologous α-carotene bioproduction requires careful balancing of lycopene ε-cyclase (LCYe) and lycopene b-cyclase (LCYb) enzymes.^35^ The initial design did consider this in account, by placing the *Arabidopsis thaliana* LCYe (AtLCYe) in front of *A. thaliana* LCYb (AtLCYb) gene (and the RBS score calculated by the **RBS calculator**^33^ resulted in 18762 and 1495, respectively, which was convenient for our purpose). However, apparently the expression level of AtLCYb was still too high relative to AtLCYe, deducing from the β-carotene accumulation. In the 2^nd^ try of plasmid construction, several different constructs were made to further lower the AtLCYb expression (Figure 6c ***(ii), (iii)*** and ***(iv)***). In one construct, where the terminator was placed between AtLCYe and AtLCYb, the *E. coli* harboring this plasmid resulted in a sole α-carotene production (Figure 6c ***(iv)***). This kind of dynamic rearranging was possible because of the the flexibility of the SA-Clo system, where I can place any parts/genes into “slots”.

As a final example, for the construction of non-natural C_50_-zeaxanthin^27^ plasmid, the initial construct placed all the genes in one operon under an arabinose promoter (Figure 6d ***(i)***). This construct resulted in production of C_50_-β-carotene, suggesting that the CrtZ enzyme expression is not enough. In the original paper that reported C_50_-zeaxanthin production,^27^ where the amount of C_50_-β-carotene was negligible, the downstream enzymes genes (*crtI_N304P_, crtY, crtZ*) were placed on a high-copy vector (pMB1) while the upstream enzymes (*fds_81,157_*and *crtM_AAS_*) were placed on a middle-copy vector (p15A). This suggests that increasing the relative expression level of *crtZ* compared to the upstream enzymes may result in C_50_-zeaxanthin production. To build a revised construct, pACm-C_50_phytoene-Lv2 plasmid (**Supplementary Table 1**) was constructed as a destination vector, and several constructs were designed aiming to maximize the relative *crtZ* expression compared to other genes. As a result, the three constructs with increased *crtZ* expression levels led to an efficient C_50_-zeaxanthin production (Figure 6d ***(iii), (iv)*** and ***(v)***). The conventional method could have required multiple rounds of tailored design and optimization, and the cloning/assembly would have been cumbersome especially when building a single-plasmid construct. Since the assembly success rate of SA-Clo is significantly high, the colony from assembly transformant could be directly used for the charactarization of carotenoid production when screening several different constructs. This example again demonstrates the relative ease for the carotenoid pathway plasmid construction.

### 5. Discussion

Carotenoids are important and useful compound, and there are many studies that have aimed optimized the plasmid for heterologous production. If the target carotenoid-of-interest has been set, and the sole goal is to maximize this carotenoid specificity/production, the best way would be to optimize the constructs extensively by using advanced combinatorial methodologies.^36^ However, if the goal is to construct various carotenoid pathway constructs that results in a single product, it will be cumbersome to repeat the optimizations. In this study, a streamlined process for carotenoid pathway construction was implemented by applying the MoClo methodology, named SA-Clo. This system would be a quick-and-easy way to construct carotenoid pathways.

While constructing plasmids using SA-Clo, the strength of the RBS remained a crucial consideration. It is well known that the translation strength mediated by the RBS in *E. coli* is influenced not only by the Shine-Dalgarno (SD) sequence but also by the upstream and downstream regions, including the coding sequence (CDS) of the gene. This is always a key consideration in gene construct design. When creating the Level-0 gene collection in this study, I included the 15 nucleotides upstream of the CDS start codon (5’-AGGAGGATTACAAA-3’) (described in results section). This the sequences of the CDS and the upstream junction may vary, which can significantly affect translation strength. However, including the RBS as a separate Level-0 part would have made this system overly complex. I considered this an acceptable trade-off because it simplified the process. In practice, the variation in RBS strength did not cause significant issues for most constructs, which we successfully created (Figure 5). The construct that was most affected by RBS strength in this study was the one for C_50_-zeaxanthin (Figure 6d). It was clear that the expression level of CrtZ was weak, and this might have been due to the low translation initiation rate. Ultimately, we achieved our goal by including the RiboJ or a new promoter upstrem the *crtZ* gene, but it is possible that the traditional method of enhancing the RBS could have also worked.

The SA-Clo system also is capable of accomodating changes to the RBS for each gene as needed. The newly designed RBS sequence can be incorporated into the PCR primer to amplify the gene-of-interest on the Level-0 plasmid. This Level-0 PCR fragment could be directly used for the Level-1 (BsaI) assembly. Furthermore, the RBS library could be created using the same strategy.

This work focused on creating plasmids for carotenoid production that result in a single product, and we have demonstrated successful examples of this goal. However, it should be noted that this does not guarantee the production of a single product in all contexts, as it depends on the *E. coli* strain and culture conditions. For instance, the constructed astaxanthin pathway produced only astaxanthin at both 24 and 48 hours when cultured in a 48 deep-well plate. In contrast, when cultured in 5 mL in 50-mL Falcon tubes, intermediates accumulated at 24 hours, and astaxanthin was produced at 48 hours (**Supplementary** Figure 3). This variability is common in metabolic and pathway engineering studies, requiring optimization for each experimental setup. Nevertheless, our system allows for quick optimization of new constructs, even under such varying conditions.

The SA-Clo system can be extended beyond carotenoids to other genetic pathways and systems. For constructs requiring the repeated combination of the same genes in various patterns, once the genes and parts are prepared in Level-0 vectors, this system facilitates the easy creation of plasmid constructs. In the future, I would like to use this sytem to create plasmids for a variety of minor carotenoids, with the aim of advancing the study of diverse carotenoid-utilizing biological systems.

## Methods

### Strains, media and reagents

*E. coli* strain used in this study were either NEB Turbo (New England BioLabs, Ipswich, MA) or DH5α (SMOBIO Technology, Inc., Taiwan). *E. coli* was cultured in LB-Lennox (Nacalai Tesque, Inc., Kyoto, #20066-24) or Terrific Broth (TB) media. Antibiotics used were chloramphenicol (FUJIFILM Wako, Osaka, #036-10571) or carbenicillin (Wako #594-02193) with the final concentration of 30 or 50 µg/mL, respectively.

### Plasmid construction

Plasmids used in this study are listed in **Supplementary Table 1**. Vectors named pACm were constructed based on pACT vector^29^ by inserting a mutation to eliminate the BsmBI restriction site (sequence provided in **Supplementary Table 2**). pAKm vector was constructed by replacing the chloramphenicol cassette with kanamycin cassette (sequence provided in in **Supplementary Table 2**). pACm-P_J23115_-lyco, pACm-P_J23115_-beta, pACm-P_J23115_-zea, pACm-P_J23115_-cantha, pACm-P_J23115_-asta (for brevity, these plasmids are referred to as pACT-P_J23115_-{zea/beta/lyco/cantha/asta}) were constructed as follows. First, pACT-P_J23107_-zea-crtZ_B_ from previous study^29^ was used as a template (renamed pACT-P_J23107_-zea) in this study. The promoter region was digested with NheI/XhoI and replaced with an oligo (5’-<u>gctagc</u>atagt**gagacc**attat**ggtctc**t<u>ctcgag</u>-3’; NheI/XhoI underlined and BsaI highlighted) to create pACT-BsaI-zea. pACT-BsaI-beta was created by removing the *crtZ* gene in pACT-BsaI-zea by digestion with KpnI/NotI. pACT-BsaI-lyco was created by removing the *crtZ-crtY* genes by digestion with KpnI/SbfI. pACT-BsaI-cantha was created by replacing *crtZ* by inserting *Brevundimonas* sp. SD212 *crtW* codon-optimized sequence^27^ into KpnI/NotI site in pACT-BsaI-zea. pACT-BsaI-asta was created by inserting *crtW* into KpnI site in pACT-BsaI-zea. Into the BsaI site of the above-constructed plasmids pACT-BsaI-{zea/beta/lyco/cantha/asta} (5 plasmids shortened for breivity), the promoter sequences (P_J23101_, P_J23107_, P_J23115_, P_J23117_) were inserted by preparing the insert DNA by annealing two oligo sets to make sticy-ends (oMF113∼oMF137 in **Supplementary Table 5**) and inserting to the vector using BsaI GoldenGate Assembly. This resulted in plasmids with different promoters, pACT-{Pxxx}-{zea/beta/lyco/cantha/asta}, where Pxxx indicates 4 different promoters. The plasmids pACm-P_J23115_-{zea/beta/lyco/cantha/asta} was constructed by removing the two BsmBI sites found in CmR casette (see pACm vector sequence in **Supplementary Table 2**) of pACT-P_J23115_-{zea/beta/lyco/cantha/asta}.

*Level-0 vectors.* The vector backbone with pMB1 ori and kanamycin resistance was prepared by amplifying the DNA fragment from pEX-K4J2 vector from Eurofin Genomics using oMF206/238 primers. The insert (GFP expression cassette flanked with Level-0 BsaI junctions) were amplified using oMF212/239 primers (see **Supplementary Table 5** for primers) and pUClac-P_tac_-sfgfp (ref) as a template. The above two DNA fragments were assembled using NEBuilder HiFi DNA assembly to obtain the pLv0 vector. Later, it became apparent that some genes could not be stably cloned into pLv0 vector because of possible leaky expression by a pseudo-promoter upstream. A terminator sequence (rrnB T_1_) was added upstream the BsaI site in pLv0 by using oMF273/274 to amplify the pLv0 vector and oMF275/276 to amplify the terminator, and by assembling these two fragments, ptLv0 vector was constructed.

*Level-1 vectors.* pUClac-P_tac_-sfgfp (ref) was used as a vector backbone. This vector was cleaved using XhoI/HindIII, and a gene fragment (gblock #103986008, **Supplementary Table 5**) of P_J23101_-lacZ cassette flanked by Level-1.1 sequence (BsaI and BsmBI) which was ordered to IDT DNA was inserted using homology assembly. Other Level-1 plasimids (Level-1.2 to 1.9 and 1.2e to 1.10e) were constructed by PCR-amplifying the P_J23101_-lacZ cassette by using primers containing each BsaI-scars described in Figure 4.

*Level-2 vectors (destination vectors).* P_tac_-sfgfp cassette was amplified using pUClac-P_tac_-sfgfp as a template using a PCR primer with BsmBI sites, and this was inserted into pACm or pAKm vector, resulting in pACm-Lv2 or pAKm-Lv2 (sequence provided in **Supplementary Table 2**). Carotenoid destination vectors (pACm-lyco-Lv2, pACm-beta-Lv2, pACm-zea-Lv2, pACm-cantha-Lv2, pACm-asta-Lv2, pACm-gamma-Lv2) was constructed by cleaving the carotenoid vectors (pACm-P_J23115_-lyco, pACm-P_J23115_-beta, pACm-P_J23115_-zea, pACm-P_J23115_-cantha, pACm-P_J23115_-asta) by HindIII and by inserting the P_tac_-sfgfp cassette was amplified using pUClac-P_tac_-sfgfp using PCR primers with BsmBI sites (oMF367/368).

### SA-Cloning of carotenoid genes: Level-1 assembly

Each Level-1 vectors (approx. 50 ng in 0.5∼1 µL) and the accompanying genes (approx. 50 ng in 0.5∼1 µL) were mixed with 0.5 µL of NEBridge Golden Gate Assembly Kit (BsaI-HF v2) reaction mix (New England Biolabs E1601), 1 µL of T4 DNA ligase buffer, 6.5 µL nuclease free water in total 10 µL reaction. This was incubated at 37°C for 5 min or 1 hour and 60°C for 5 min, following the manufacture’s guideline. The 1 µL aliquot was transformed into 10 µL of NEB Turbo or DH5α competent cells, 100-200 µL of SOC media was added and incubated for 1 hour. A portion (20 µL) was plated on agar plate supplemented with X-gal, IPTG and antibiotics, and another portion of 1-2 µL was inoculated into 2-mL LB with antibiotics and shaken at 37C, 200 rpm overnight. After confirming the cfu on the ager plate, plasmids were purified from the 2-mL culture and eluted using 50 µL EB buffer from QIAGEN. The concentration usually was 50 ng/µL. This was used immediately for the Level-2 assembly.

### SA-Cloning of carotenoid genes: Level-2 assembly

The prepared Level-1 parts (approx. 25 ng in 0.5 µL) and Level-2 vector (approx. 25 ng in 0.5 µL) were added to an 8-strip PCR tube. These DNA were mixed with 1 µL of T4 DNA ligase buffer (New England Biolabs), 1 µL of NEBridge Golden Gate Assembly Kit (BsmBI-v2) reaction mix (New England Biolabs E1602), nuclease free water up to 10 µL. The mixture was placed in thermal cycler and ran with the following protocol: (42°C, 1 min ➔ 16°C, 1 min) x 40-60 cycles ➔ 60°C, 5 min. Using the NEBridge Golden Gate Assembly Kit (rather than preparing your own enzyme mix) significantly increased the assembly efficiency.

### *E. coli* carotenoid production and extraction

*E. coli* NEB Turbo or DH5α was transformed using the plasmids and plated on an LB-agar plate supplemented with antibiotics. The single colonies were picked and inoculated into a 2-mL LB-media with antibiotics in a 2-position round bottom 15-mL tube (EVERGREEN, #222-2094-050) and incubated at 37°C, 200 rpm overnight (approx. 16 hours). An aliquot (100 µL) of these pre-culture cells were inoculated into a 5 or 10-mL TB media supplemented with antibiotics in 50-mL falcon tube with a filter cap (Greiner Bio-One #227245) and cultured at 30°C or 25°C at 200 rpm for 48 hours using a TAITEC BR-43FL incubator shaker.

The cells were harvested at 7000g for 3 min, and the cell pellet were washed with saline and harvested again. The cell pellet was vortexed for 1 min to soften. Acetone 5 mL were added to the softened-pellet and vortexed immediately for 3 min. The tubes were centrifuged at max speed for 3 min and the acetone layer were collected into a new 50-mL tube. For natural (C_40_) carotenoids, this acetone extract was used directly for LC analysis.

For the non-natural (C_50_) carotenoids, the carotenoid extract was further concentrated as follows. To the acetone-extract in 50-mL tube, 1-mL of hexane and 30-mL of 10% NaCl were added, followed by centrifugation at 7000g for 3 min. The top hexane layer was collected into a new 4-mL glass vial, followed by evaporation using Smart Evaporator™ C4-Lite (BioChromato, Fujisawa, Japan). The dried carotenoids were resuspended in a 100 µL of mixed solution of THF:methanol of 1:1, and this was diluted 1/10 into methanol to use for the LC analysis.

### LC-PDA-MS analysis of extracted carotenoids

The equipment used were Waters UPLC H-Class PLUS consisting of a quaternary pump with degasser, autosampler and photodiode array (PDA) in combination with a Xevo TQ-S micro triple quadrupole mass spectrometer. The column used was Waters ACQUITY UPLC BEH C18 1.7 µm column (2.1 mm x 100 mm) equipped with a C18 VanGuard Pre-column. The mobile phase used were A: water/acetonitrile = 10/90, B: acetonitrile, C: methanol, D: isopropanol with an elution flux of 0.2 ml/min with the following gradient program (total 15 min). Starting from [100% A], 0-2 min gradient to [70% A, 30% B]; 2-4 min gradient to [35% B, 15% C, 50% D] and hold for 8 min until 12 min; switch to [100% A] and hold for 3 min. The elution was monitored with absorbance at 450 nm. The data was analyzed using MassLynx software.

### Quantification of carotenoid produced by E. coli in Fig. 2b

*E. coli* strain was transformed by the indicated plasmids, plated to LB agar plates with appropriate antibiotics, and incubated overnight at 37°C. Single colonies were picked to inoculate 500 µL of LB broth with appropriate antibiotics in a 96-well deep well plate. The cells were cultured using an incubator shaker (TAITEC MBR-034P) at 1000 rpm, 37°C for 16 hours. A portion (20 µL) of this overnight culture was transferred to a fresh 2 mL TB medium with antibiotics in a 48-well deep well plate. The plate was shaken at 1000 rpm, 30°C for 48 hours.

One milliliter of liquid culture was collected into a 2-mL collection tube and centrifuged at 8000x*g* for 1 min. The supernatant was removed, the pellet was resuspended in 1% NaClaq, and the resuspension was centrifuged again at the same speed, and the cells were vortexed briefly to loosen the pellet. One milliliter of acetone was added to the cell pellet and vortexed immediately for 1 min. The collection tube was centrifuged for 2 min at max speed, and the carotenoid-containing extract supernatant was collected into a polypropylene 48-well deep well plate. One-hundred microliters of the extract was transferred to a polypropylene 96-well plate and the absorbance spectrum were measured using Tekan Spark microplate reader. Carotenoids were quantified by calculating the concentration using Beer–Lambert law using the specific absorbance coefficient A (1%, 1cm) (lycopene 3446, β-carotene 2500, zeaxanthin 2340, canthaxanthin 2200, astaxanthin 2100).

## Data availability statement

All data generated or analysed during this study are included in this published article and its Supplementary Information files.

## Acknowledgements

M.F. thanks M. Nariai., M. Komatsu and M. Hayakawa for their assistance. This work was supported by JSPS KAKENHI Grant Numbers 19K23670, 20K15442, 23K26837, 23H02144.

## Additional Information

Competing Interests Statement: The author declare no competing interests.

