## Supplementary File for "Systematic Plasmid Engineering for Targeted Carotenoid Synthesis in Bacteria"

Maiko Furubayashi

Bioproduction Research Institute (BPRI), National Institute of Advanced Industrial Science and Technology (AIST), Sapporo, Hokkaido, Japan

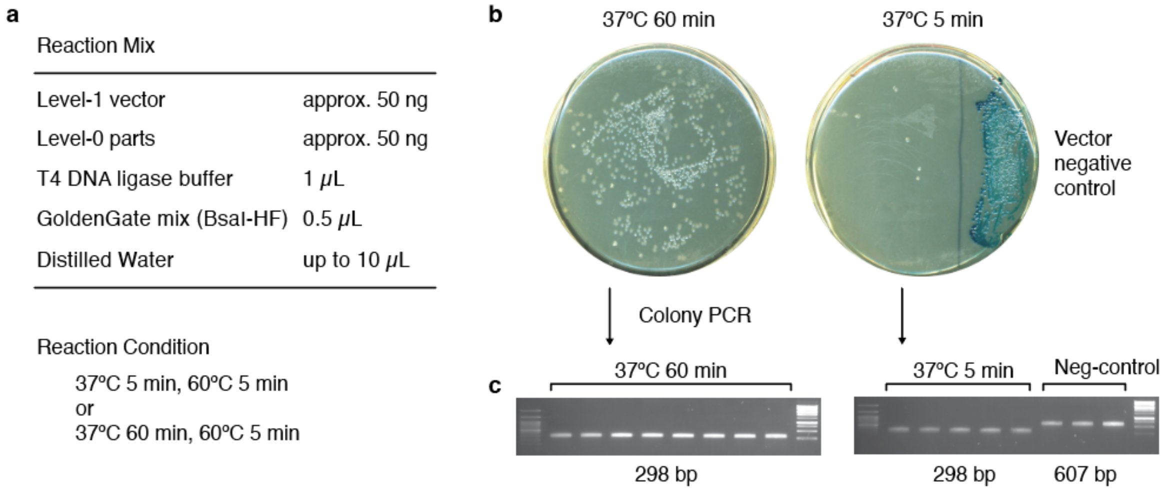


**Supplementary Figure 1. Level-1 assembly.**
(**a**) Reaction condition (see Methods). The 37ºC incubation was performed either 5 or 60 min. One microliter of the reaction mixture was used to transform 10 μL of competent cells. (**b**) Representative agar plates of *E. coli* transformants of Level-1 assembly (1/10 of the transformants plated on agar). Vector control (with lacZ) are plated as a control, showing blue color. (**c**) Agarose gel electrophoresis of colony PCR. The correct fragment length is shown on the bottom (298 bp for positive clones, 607 bp for negative clones).

a

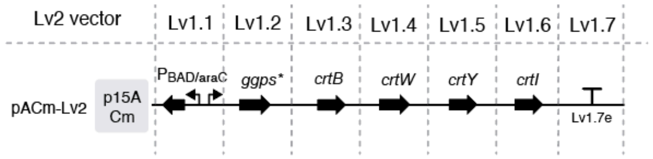


a

| Reaction Mix | |
|---|---|
| Level-2 vector | approx. 50 ng |
| Level-1 genes/parts | approx. 50 ng each |
| T4 DNA ligase buffer | 1 $\mu$L |
| GoldenGate mix (BsmBI-HF) | 1 $\mu$L |
| Distilled Water | up to 10 $\mu$L |

Reaction Condition

[ 37°C 1 min, 16°C 1 min ] x 40-60 cycles, 60°C 5 min

c

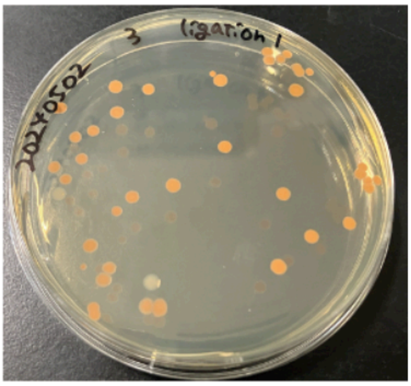

**Supplementary Figure 2. Level-2 assembly.**

(**a**) Assembly constructs. (**b**) Reaction condition (see Methods). (**c**) Representative agar plates of E. coli transformants of Level-2 assembly. No background (GFP) coloneis were observed. Orange color indicates correct assembly (canthaxanthin biosynthesis).

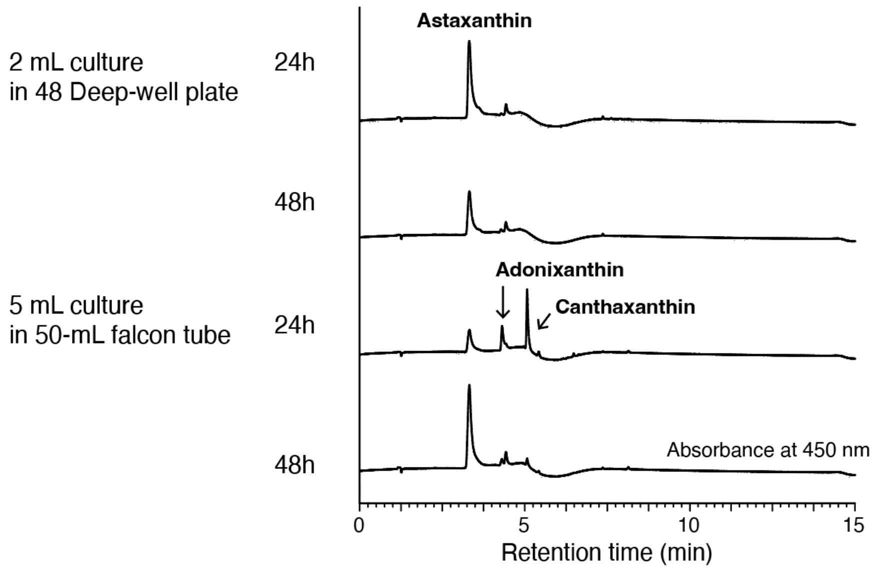


**Supplementary Figure 3. Astaxanthin production in different conditions.** The astaxanthin plasmid used in Fig. 2a-b are used in this figure. The experimental condition for 5-mL culture are described in methods under "*E. coli* carotenoid production and extraction". For the 2-mL culture, see "Quantification of carotenoid produced by E. coli in Fig. 2b" in methods.

### Supplementary Table 1. Plasmids used in this study.[1]

| Plasmid name[2] | Description[3] | Other names |
|---|---|---|
| **Vector** | | |
| pACT | p15A, CmR; ref 29 in main text | |
| pACm | p15A, CmR; deleted BsmBI from pACT vector | |
| pAKT | p15A, KmR; replaced CmR with KmR in pACT vector | |
| pAKm | P15A, KmR; eliminated pseudo promoter in pAKT | |
| **Carotenoid plasmids on p15A** | | |
| pACT-P[xxx]-lyco | $P_{J23xxx}$-ggps*$_{Gs}$, crtB$_{Pa}$, crtI$_{Pa}$ | pMF559, 561, 562 |
| pACT-P[xxx]-beta | $P_{J23xxx}$-ggps*$_{Gs}$, crtB$_{Pa}$, crtI$_{Pa}$, crtY$_{Pa}$ | pMF563, 565, 566 |
| pACT-P[xxx]-zea | $P_{J23xxx}$-ggps*$_{Gs}$, crtB$_{Pa}$, crtI$_{Pa}$, crtY$_{Pa}$, crtZ$_{Br}$ | pMF547, 548,549 |
| pACT-P[xxx]-can | $P_{J23xxx}$-ggps*$_{Gs}$, crtB$_{Pa}$, crtI$_{Pa}$, crtY$_{Pa}$, crtW$_{Br}$ | pMF567, 569, 570 |
| pACT-P[xxx]-asta | $P_{J23xxx}$-ggps*$_{Gs}$, crtB$_{Pa}$, crtI$_{Pa}$, crtY$_{Pa}$, crtZ$_{Br}$, crtW$_{Br}$ | pMF571, 573, 574 |
| pACm-$P_{J23115}$-lyco | $P_{J23115}$-ggps*$_{Gs}$, crtB$_{Pa}$, crtI$_{Pa}$ | pMF1074 |
| pACm-$P_{J23115}$-beta | $P_{J23115}$-ggps*$_{Gs}$, crtB$_{Pa}$, crtI$_{Pa}$, crtY$_{Pa}$ | pMF910 |
| pACm-$P_{J23115}$-zea | $P_{J23115}$-ggps*$_{Gs}$, crtB$_{Pa}$, crtI$_{Pa}$, crtY$_{Pa}$, crtZ$_{Br}$ | pMF911 |
| pACm-$P_{J23115}$-can | $P_{J23115}$-ggps*$_{Gs}$, crtB$_{Pa}$, crtI$_{Pa}$, crtY$_{Pa}$, crtW$_{Br}$ | pMF912 |
| pACm-$P_{J23115}$-asta | $P_{J23115}$-ggps*$_{Gs}$, crtB$_{Pa}$, crtI$_{Pa}$, crtY$_{Pa}$, crtZ$_{Br}$, crtW$_{Br}$ | pMF867 |
| **Level-0 vectors** | | |
| pLv0 | pUC, $Kan^R$ | pMF710 |
| ptLv0 | Terminator inserted upstream the Level-0 cloning site in pLv0 to prevent read-through | pMF764 |
| **Level-1 vectors** | | |
| pLv1.1, 1.2 … 1.9 | Ptac/lacI promoter upstream the Level-1 cloning site | pMF668,799,800,801,844,1072, 1182, 1183, 1184 |
| pLv1.1e, 1.2e, … 1.10e | Ptac/lacI promoter upstream the Level-1 cloning site | pMF756,757,802,667,803,845, 1073,1185, 1186, 1187 |
| ptLv1.1, 1.2 … 1.8 | Terminator upstream the Level-1 cloning site | pMF1144~1150 |
| ptLv1.1e, 1.2e … 1.9e | Terminator upstream the Level-1 cloning site | pMF1151~1157 |
| **Level-2 vectors** | | |
| pACm-Lv2 | p15A, CmR, Lv2-casette | pMF939 |
| pAKm-Lv2 | p15A, KmR, Lv2-casette | |
| pACm-lyco-Lv2 | $P_{J23115}$-ggps*$_{Gs}$, crtB$_{Pa}$, crtI$_{Pa}$, Lv2-casette | pMF1074 |
| pACm-beta-Lv2 | $P_{J23115}$-ggps*$_{Gs}$, crtB$_{Pa}$, crtI$_{Pa}$, crtY$_{Pa}$, Lv2-casette | pMF910 |
| pACm-zea-Lv2 | $P_{J23115}$-ggps*$_{Gs}$, crtB$_{Pa}$, crtI$_{Pa}$, crtY$_{Pa}$, crtZ$_{Br}$, Lv2-casette | pMF911 |
| pACm-can-Lv2 | $P_{J23115}$-ggps*$_{Gs}$, crtB$_{Pa}$, crtI$_{Pa}$, crtY$_{Pa}$, crtW$_{Br}$, Lv2-casette | pMF912 |
| pACm-asta-Lv2 | $P_{J23115}$-ggps*$_{Gs}$, crtB$_{Pa}$, crtI$_{Pa}$, crtY$_{Pa}$, crtZ$_{Br}$, crtW$_{Br}$, Lv2-casette | pMF867 |
| pACm-C50phy-Lv2 | $P_{J23117}$-ggps$_{Y81A,V157A_Gs}$, crtM$_{AAS_Sa}$, Lv2-casette | pMF1110 |

[1]The plasmids constructed by SA-Cloning are not shown. See **Supplementary Table 4**.
[2]P[xxx]: $P_{J23101}$, $P_{J23107}$, $P_{J23117}$
[3]Subscript stands for; Gs: *G. stearothermophillus*, Pa: *P. ananatis*; Bl: *B. linens*; Br: *Brevundimonas* sp. SD212

### Supplementary Table 2. Sequences for Level-0, 1 and 2 vectors.

| Name | Sequence |
|---|---|
| pLv0<br><br>BsaI:<br>recognition<br>**sticky-end** | GGTCTCT**CTAT**tgaggaggattacatatgaattcagctgttgacaattaatcatcggctcgtataatgtgtggaattgtgagcggataacaatttcacacaggaaacactcgaggat<br>taaataaggaggaataaaccatgggtcatcaccaccatcatcacggtggcgcgtctaaaggcgaagaactgttcaccggcgtagttccgatcctggttgaactggacggtgacgtta<br>atggtcataagttctctgttcgtggtgaaggtgagggcgacgcgaccaacggtaaactgaccctgaaattcatctgcaccactggcaaactgccggttccgtggccgactctggttac<br>caccctgacctatggtgttcagtgcttctctcgttacccggatcacatgaaacagcacgacttcttcaaatctgcgatgccggagggttatgttcaggaacgtaccatctctttcaagg<br>atgacggcacctacaaaacccgtgccgaagttaaattcgagggtgatacgctggtaaaccgcatcgaactgaaaggtatcgacttcaaagaggacggtaatatcctcggtcacaa<br>gctggaatacaacttcaactctcacaacgtttacatcaccgcggacaaacagaaaaacggtatcaaagcgaactttaagatccgtcacaatgttgaagacggcagcgttcagctc<br>gctgaccactaccaacaaaataccccgattggcgacggtccggttctgctgccggacaaccactatctgtctacccagtctgcgctctctaaggacccgaacgagaaacgtgacc<br>acatggtgctgctggagttcgtgaccgcagcgggcatcacgcacggcatggacgaactgtacaaataataaaagcttgccaggcatcaaata**TGTT**TGAGACCagtcagtgga<br>tacgccaaatgccttcatgcgcccaaaacatgttggcatgttcacgatgtgcccaacgatgtccttcccaacgtgctctaccggcatgatcaaagctatgcgccttaaacaggtcat<br>gcgctgaatagtggcgttcgattgcccaaccgtagaagtctttgcctgtcgtgccagctgcattaatgaatcggccaacgcgcggggagaggcggtttgcgtattgggcagagaatat<br>aaaaagccagattattaatccggctttttattatttgctcttccgcttcctcgctcactgactcgctgcgctcggtcgttcggctgcggcgagcggtatcagctcactcaaaggcggtaa<br>tacggttatccacagaatcaggggataacgcaggaaagaacatgtgagcaaaaggccagcaaaaggccaggaaccgtaaaaaggccgcgttgctggcgtttttccataggctcc<br>gcccccctgacgagcatcacaaaaatcgacgctcaagtcagaggtggcgaaacccgacaggactataaagataccaggcgtttccccctggaagctccctcgtgcgctctcctg<br>ttccgaccctgccgcttaccggatacctgtccgcctttctcccttcgggaagcgtggcgctttctcatagctcacgctgtaggtatctcagttcggtgtaggtcgttcgctccaagctggg<br>ctgtgtgcacgaaccccccgttcagcccgaccgctgcgccttatccggtaactatcgtcttgagtccaacccggtaagacacgacttatcgccactggcagcagccactggtaac<br>aggattagcagagcgaggtatgtaggcggtgctacagagttcttgaagtggtggcctaactacggctacactagaagaacagtatttggtatctgcgctctgctgaagccagttacctt<br>cggaaaaagagttggtagctcttgatccggcaaacaaaccaccgctggtagcggtggtttttttgtttgcaagcagcagattacgcgcagaaaaaaaggatctcaagaagatcctttg<br>atcttttctacggggtctgacgctcagtggaacgaaaactcacgttaagggattttggtcatgagattatcaaaaaggatcttcacctagatccttttaaattaaaaatgaagttttaaat<br>caatctaaagtatatatgagtaaaaatattccggaattgccagctggggcgccctctggtaaggttgggaagccctgcaaagtaaactggatggctttcttgccgccaaggatctgatg<br>gcgcaggggatcaagatctgatcaagagacaggatgaggatcgtttcgcatgattgaacaagatggattgcacgcaggttctccggccgcttgggtggagaggctattcggctatgac<br>tgggcaCAACAGACAATcggctgctctgatgccgccgtgttccggctgtcagcgcaggggcgcccggttctttttgtcaagaccgacctgtccggtgccctgaatgaactgcagg<br>acgaggcagcgcggctatcgtggctggccacgacgggcgttccttgcgcagctgtgctcgacgttgtcactgaagcgggaagggactggctgctattgggcgaagtgccggggcag<br>gatctcctgtcatcccaccttgctcctgccgagaaagtatccatcatggctgatgcaatgcggcggctgcatacgcttgatccggctacctgcccattcgaccaccaagcgaaacat<br>cgcatcgagcgagcacgtactcggatggaagccggtcttgtcgatcaggatgatctggacgaagagcatcaggggctcgcgccagccgaactgttcgccaggctcaaggcgcgca<br>tgcccgacggcgaggatctcgtcgtgacccatggcgatgcctgcttgccgaatatcatggtggaaaatggccgcttttctggattcatcgactgtggccggctgggtgtggcggaccgc<br>tatcaggacatagcgttggctacccgtgatattgctgaagagcttggcggcgaatgggctgaccgcttcctcgtgctttacggtatcgccgctcccgattcgcagcgcatcgccttctat<br>cgccttcttgacgagttcttctgaaccggtaatattattgaagcatttatcagggttattgtctcatgagcggatacatatttgaatgtatttagaaaaataaacaaataggggttccgcgc<br>acatttccccgaaaagtgccacctgacgtctaagaaaccattattatcatgacattaacctataaaaataggcgtatcacgaggcccttt |
| ptLv0<br><br>BsaI:<br>recognition<br>**sticky-end** | GGTCTCT**CTAT**tgaggaggattacatatgaattcagctgttgacaattaatcatcggctcgtataatgtgtggaattgtgagcggataacaatttcacacaggaaacactcgaggat<br>taaataaggaggaataaaccatgggtcatcaccaccatcatcacggtggcgcgtctaaaggcgaagaactgttcaccggcgtagttccgatcctggttgaactggacggtgacgtta<br>atggtcataagttctctgttcgtggtgaaggtgagggcgacgcgaccaacggtaaactgaccctgaaattcatctgcaccactggcaaactgccggttccgtggccgactctggttac<br>caccctgacctatggtgttcagtgcttctctcgttacccggatcacatgaaacagcacgacttcttcaaatctgcgatgccggagggttatgttcaggaacgtaccatctctttcaagg<br>atgacggcacctacaaaacccgtgccgaagttaaattcgagggtgatacgctggtaaaccgcatcgaactgaaaggtatcgacttcaaagaggacggtaatatcctcggtcacaa<br>gctggaatacaacttcaactctcacaacgtttacatcaccgcggacaaacagaaaaacggtatcaaagcgaactttaagatccgtcacaatgttgaagacggcagcgttcagctc<br>gctgaccactaccaacaaaataccccgattggcgacggtccggttctgctgccggacaaccactatctgtctacccagtctgcgctctctaaggacccgaacgagaaacgtgacc<br>acatggtgctgctggagttcgtgaccgcagcgggcatcacgcacggcatggacgaactgtacaaataataaaagcttgccaggcatcaaata**TGTT**TGAGACCagtcagtgga<br>tacgccaaatgccttcatgcgcccaaaacatgttggcatgttcacgatgtgcccaacgatgtccttcccaacgtgctctaccggcatgatcaaagctatgcgccttaaacaggtcat<br>gcgctgaatagtggcgttcgattgcccaaccgtagaagtctttgcctgtcgtgccagctgcattaatgaatcggccaacgcgcggggagaggcggtttgcgtattgggcagagaatat<br>aaaaagccagattattaatccggctttttattatttgctcttccgcttcctcgctcactgactcgctgcgctcggtcgttcggctgcggcgagcggtatcagctcactcaaaggcggtaa<br>tacggttatccacagaatcaggggataacgcaggaaagaacatgtgagcaaaaggccagcaaaaggccaggaaccgtaaaaaggccgcgttgctggcgtttttccataggctcc<br>gcccccctgacgagcatcacaaaaatcgacgctcaagtcagaggtggcgaaacccgacaggactataaagataccaggcgtttccccctggaagctccctcgtgcgctctcctg<br>ttccgaccctgccgcttaccggatacctgtccgcctttctcccttcgggaagcgtggcgctttctcatagctcacgctgtaggtatctcagttcggtgtaggtcgttcgctccaagctggg<br>ctgtgtgcacgaaccccccgttcagcccgaccgctgcgccttatccggtaactatcgtcttgagtccaacccggtaagacacgacttatcgccactggcagcagccactggtaac<br>aggattagcagagcgaggtatgtaggcggtgctacagagttcttgaagtggtggcctaactacggctacactagaagaacagtatttggtatctgcgctctgctgaagccagttacctt<br>cggaaaaagagttggtagctcttgatccggcaaacaaaccaccgctggtagcggtggtttttttgtttgcaagcagcagattacgcgcagaaaaaaaggatctcaagaagatcctttg<br>atcttttctacggggtctgacgctcagtggaacgaaaactcacgttaagggattttggtcatgagattatcaaaaaggatcttcacctagatccttttaaattaaaaatgaagttttaaat<br>caatctaaagtatatatgagtaaaaatattccggaattgccagctggggcgccctctggtaaggttgggaagccctgcaaagtaaactggatggctttcttgccgccaaggatctgatg<br>gcgcaggggatcaagatctgatcaagagacaggatgaggatcgtttcgcatgattgaacaagatggattgcacgcaggttctccggccgcttgggtggagaggctattcggctatgac<br>tgggcacaacagacaatcggctgctctgatgccgccgtgttccggctgtcagcgcaggggcgcccggttctttttgtcaagaccgacctgtccggtgccctgaatgaactgcaggac<br>gaggcagcgcggctatcgtggctggccacgacgggcgttccttgcgcagctgtgctcgacgttgtcactgaagcgggaagggactggctgctattgggcgaagtgccggggcaggat<br>ctcctgtcatcccaccttgctcctgccgagaaagtatccatcatggctgatgcaatgcggcggctgcatacgcttgatccggctacctgcccattcgaccaccaagcgaaacatcg<br>catcgagcgagcacgtactcggatggaagccggtcttgtcgatcaggatgatctggacgaagagcatcaggggctcgcgccagccgaactgttcgccaggctcaaggcgcgcatg<br>cccgacggcgaggatctcgtcgtgacccatggcgatgcctgcttgccgaatatcatggtggaaaatggccgcttttctggattcatcgactgtggccggctgggtgtggcggaccgcta<br>tcaggacatagcgttggctacccgtgatattgctgaagagcttggcggcgaatgggctgaccgcttcctcgtgctttacggtatcgccgctcccgattcgcagcgcatcgccttctatc<br>gccttcttgacgagttcttctgaaccggtaatattattgaagcatttatcagggttattgtctcatgagcggatacatatttgaaatgcgagagtagggaactgCCAGGCATCAAATA<br>AAACGAAAGGCTCAGTCGAAAGACTGGGCCTTTCGTTTTATCTGTTGTTTGTCGGTGAACGCTCTCctgagtaggacaaatccgccgctgacgtctaag<br>aaaccattattatcatgacattaacctataaaaataggcgtatcacgaggcccttt |
| pLv1.x<br><br>BsaI:<br>recognition<br>**sticky-end**<br><br>BsmBI:<br>recognition<br>sticky-end | CGTCTCAXXXX**CTAT**AGAGACCttacagctagctcagtcctaggtattatgctagcgctatgaccatggtacaaagaggagaaaggacatgggtgacccgaaactgctgct<br>ggaagttccgcacgctatctcttcttctccgggtaactctctggctgttgttctgcaacgtcgtgactgggaaaacccgggtgttacccagctgaaccgtctggctgctcacccgccgtt<br>cgcttcttggcgtaactctgaagaagctcgtaccgaccgtccgtcccagcagctgcgttctctgatccgtctgctgaccaaaccggaacgtaaactgtcttggctgctgccgccgctg<br>tctaacaactaacgcaaaaaaGGTCTCA**TGTT**YYYYAGAGACGcaagcttgccaggcatcaaataaaacgaaaggctcagtcgaaagactgggcctttcgttttatctgtt<br>gtttgtcggtgaacgctctcctgagtaggacaaatccgcagacgaaagggcctcgtgatacgcctatttttataggttaatgtcatgataataatggtttcttagacgtcaggtggcacttt<br>tcggggaaatgtgcgcggaacccctatttgtttatttttctaaatacattcaaatatgtatccgctcatgagacaataaccctgataaatgcttcaataatattgaaaaaggaagagtatg<br>agtattcaacatttccgtgtcgcccttattcccttttttgcggcattttgccttcctgtttttgctcacccagaaacgctggtgaaagtaaaagatgctgaagatcagttgggtgcacgagtg<br>ggttacatcgaactggatctcaacagcggtaagatccttgagagttttcgccccgaagaacgttttccaatgatgagcacttttaaagttctgctatgtggcgcggtattatcccgtattg<br>acgccgggcaagagcaactcggtcgccgcatacactattctcagaatgacttggttgagtactcaccagtcacagaaaagcatcttacggatggcatgacagtaagagaattatgc<br>agtgctgccataaccatgagtgataacactgcggccaacttacttctgacaacgatcggaggaccgaaggagctaaccgcttttttgcacaacatgggggatcatgtaactcgcctt<br>gatcgttgggaaccggagctgaatgaagccataccaaacgacgagcgtgacaccacgatgcctgtagcaatggcaacaacgttgcgcaaactattaactggcgaactacttactc<br>tagcttcccggcaacaattaatagactggatggaggcggataaagttgcaggaccacttctgcgctcggcccttccggctggctggtttattgctgataaatctggagccggtgagcgt<br>gggtcacgcggtatcattgcagcactggggccagatggtaagccctcccgtatcgtagttatctacacgacggggagtcaggcaactatggatgaacgaaatagacagatcgctga<br>gataggtgcctcactgattaagcattggtaactgtcagaccaagtttactcatatatactttagattgatttaaaacttcatttttaatttaaaaggatctaggtgaagatcctttttgataat<br>ctcatgaccaaaatcccttaacgtgagttttcgttccactgagcgtcagaccccgtagaaaagatcaaaggatcttcttgagatcctttttttctgcgcgtaatctgctgcttgcaaaca<br>aaaaaaccaccgctaccagcggtggtttgtttgccggatcaagagctaccaactctttttccgaaggtaactggcttcagcagagcgcagataccaaatactgtccttctagtgtagc<br>cgtagttaggccaccacttcaagaactctgtagcaccgcctacatacctcgctctgctaatcctgttaccagtggctgctgccagtggcgataagtcgtgtcttaccgggttggactca<br>agacgatagttaccggataaggcgcagcggtcgggctgaacggggggttcgtgcacacagcccagcttggagcgaacgacctacaccgaactgagatacctacagcgtgagcta<br>tgagaaagcgccacgcttcccgaagggagaaaggcggacaggtatccggtaagcggcagggtcggaacaggagagcgcacgagggagcttccagggggaaacgcctggtatct |

| | |
|---|---|
| | ttatagtcctgtcgggtttcgccacctctgacttgagcgtcgatttttgtgatgctcgtcaggggggcggagcctatggaaaaacgccagcaacgcggcctttttacggttcctggcctttt<br>gctggccttttgctcacatgttctttcctgcgttatcccctgattctgtggataaccgtattaccgcctttgagtgagctgataccgctcgccgcagccgaacgaccgagcgcagcgagt<br>cagtgagcgaggaagcggaagagtgcctaatgagtgaactcacattaattgcgttgcgctcactgcccgctttccagtcgggaaacctgtcgtgccagctgcattaatgaatcggcc<br>aacgcgcggggagaggcggtttgcgtattgggcgccagggtggtttttcttttcaccagtgaaacgggcaacagctgattgcccttcaccgcctggccctgagagagttgcagcaagc<br>ggtccacgctggtttgccccagcaggcgaaaatcctgtttgatggtggttaacggcgggatataacatgagctgtcttcggtatcgtcgtatcccactaccgagatatccgcaccaac<br>gcgcagcccggactcggtaatggcgcgcattgcgcccagcgccatctgatcgttggcaaccagcatcgcagtgggaacgatgccctcattcagcatttgcatggtttgttgaaaacc<br>ggacatggcactccagtcgccttcccgttccgctatcggctgaatttgattgcgagtgagatatttatgccagccagccagacgcagacgcgccgagacagaacttaatgggcccgc<br>taacagcgcgatttgctggtgacccaatgcgaccagatgctccacgcccagtcgcgtaccgtcttcatgggagaaaataatactgttgatgggtgtctggtcagagacatcaagaaa<br>taacgccggaacattagtgcaggcagcttccacagcaatggcatcctggtcatccagcggatagttaatgatcagcccactgacgcgttgcgcgagaagattgtgcaccgccgcttt<br>acaggcttcgacgccgcttcgttctaccatcgacaccaccacgctggcacccagttgatcggcgcgagatttaatcgccgcgacaatttgcgacggcgcgtgcagggccagactg<br>gaggtggcaacgccaatcagcaacgactgtttgcccgccagttgttgtgccacgcggttgggaatgtaattcagctccgccatcgccgcttccactttttcccgcgttttcgcagaaac<br>gtggctggcctggttcaccacgcgggaaacggtctgataagagacaccggcatactctgcgacatcgtataacgttactggtttcacattcaccaccctgaattgactctcttccggg<br>cgctatcatgccataccgcgaaaggttttgcaccattcgatgggaattcagctgttgacaattaatcatcggctcgtataatgtgtggaattgtgagcggataacaatttcacacaggaa<br>acactcgagta |
| ptLv1.x<br><br>BsaI:<br>recognition<br>**sticky-end**<br><br>BsmBI:<br>recognition<br>sticky-end | CGTCTCAXXXX**CTAT**AGAGACCttacagctagctcagtcctaggtattatgctagcgctatgaccatggtacaaagaggagaaaggacatgggtgacccgaaactgctgct<br>ggaagttccgcacgctatctcttcttctccgggtaactctctggctgttgttctgcaacgtcgtgactgggaaaacccgggtgttacccagctgaaccgtctggctgctcacccgccgtt<br>cgcttcttggcgtaactctgaagaagctcgtaccgaccgtccgtcccagcagctgcgttctctgatccgtctgctgaccaaaccggaacgtaaactgtcttggctgctgccgccgctg<br>tctaacaactaacgcaaaaaaGGTCTCA**TGTT**YYYYAGAGACcaagcttgccaggcatcaaataaaacgaaaggctcagtcgaaagactgggcctttcgttttatctgttgt<br>ttgtcggtgaacgctctcctgagtaggacaaatccgcagacgaaagggcctcgtgatacgcctatttttataggttaatgtcatgataataatggtttcttagacgtcaggtggcacttttc<br>ggggaaatgtgcgcggaacccctatttgtttatttttctaaatacattcaaatatgtatccgctcatgagacaataaccctgataaatgcttcaataatattgaaaaaggaagagtatga<br>gtattcaacatttccgtgtcgcccttattcccttttttgcggcattttgccttcctgtttttgctcacccagaaacgctggtgaaagtaaaagatgctgaagatcagttgggtgcacgagtgg<br>gttacatcgaactggatctcaacagcggtaagatccttgagagttttcgccccgaagaacgttttccaatgatgagcacttttaaagttctgctatgtggcgcggtattatcccgtattga<br>cgccgggcaagagcaactcggtcgccgcatacactattctcagaatgacttggttgagtactcaccagtcacagaaaagcatcttacggatggcatgacagtaagagaattatgca<br>gtgctgccataaccatgagtgataacactgcggccaacttacttctgacaacgatcggaggaccgaaggagctaaccgcttttttgcacaacatgggggatcatgtaactcgccttg<br>atcgttgggaaccggagctgaatgaagccataccaaacgacgagcgtgacaccacgatgcctgtagcaatggcaacaacgttgcgcaaactattaactggcgaactacttactct<br>agcttcccggcaacaattaatagactggatggaggcggataaagttgcaggaccacttctgcgctcggcccttccggctggctggtttattgctgataaatctggagccggtgagcgtg<br>ggtcacgcggtatcattgcagcactggggccagatggtaagccctcccgtatcgtagttatctacacgacggggagtcaggcaactatggatgaacgaaatagacagatcgctgag<br>ataggtgcctcactgattaagcattggtaactgtcagaccaagtttactcatatatactttagattgatttaaaacttcatttttaatttaaaaggatctaggtgaagatcctttttgataatct<br>catgaccaaaatcccttaacgtgagttttcgttccactgagcgtcagaccccgtagaaaagatcaaaggatcttcttgagatcctttttttctgcgcgtaatctgctgcttgcaaacaaa<br>aaaaccaccgctaccagcggtggtttgtttgccggatcaagagctaccaactctttttccgaaggtaactggcttcagcagagcgcagataccaaatactgtccttctagtgtagccg<br>tagttaggccaccacttcaagaactctgtagcaccgcctacatacctcgctctgctaatcctgttaccagtggctgctgccagtggcgataagtcgtgtcttaccgggttggactcaag<br>acgatagttaccggataaggcgcagcggtcgggctgaacggggggttcgtgcacacagcccagcttggagcgaacgacctacaccgaactgagatacctacagcgtgagctatg<br>agaaagcgccacgcttcccgaagggagaaaggcggacaggtatccggtaagcggcagggtcggaacaggagagcgcacgagggagcttccagggggaaacgcctggtatcttt<br>atagtcctgtcgggtttcgccacctctgacttgagcgtcgatttttgtgatgctcgtcaggggggcggagcctatggaaaaacgccagcaacgcggcctttttacggttcctggccttttg<br>ctggccttttgctcacatgttctttcctgcgttatcccctgattctgtggataaccgtattaccgcctttgagtgagctgataccgctcgccgcagccgaacgaccgagcgcagcgagtc<br>agtgagcgaggaagcggaagaggccacatgaagcacttctcaattgttcagaacgctcggtcttgcacaccgggcgtttttttctttagcacaatttcacacaggaaacactcgagta |
| pACm-Lv2<br><br>BsmBI:<br>recognition<br>sticky-end | caccggtgcagcctttttttctcctgccacatgaagcacttctcaattgttcagaacgctcggtcttgcacaccgggcgtttttttctttagcATCTTGAGACGctgttgacaattaatc<br>atcggctcgtataatgtgtggaattgtgagcggataacaatttcacacaggaaacactcgaggattaaataaggaggaataaaccatgggtcatcaccaccatcatcacggtggcgc<br>gtctaaaggcgaagaactgttcaccggcgtagttccgatcctggttgaactggacggtgacgttaatggtcataagttctctgttcgtggtgaaggtgagggcgacgcgaccaacggta<br>aactgaccctgaaattcatctgcaccactggcaaactgccggttccgtggccgactctggttaccaccctgacctatggtgttcagtgcttctctcgttacccggatcacatgaaaca<br>gcacgacttcttcaaatctgcgatgccggagggttatgttcaggaacgtaccatctctttcaaggatgacggcacctacaaaacccgtgccgaagttaaattcgagggtgatacgctg<br>gtaaaccgcatcgaactgaaaggtatcgacttcaaagaggacggtaatatcctcggtcacaagctggaatacaacttcaactctcacaacgtttacatcaccgcggacaaacaga<br>aaaacggtatcaaagcgaactttaagatccgtcacaatgttgaagacggcagcgttcagctcgctgaccactaccaacaaaataccccgattggcgacggtccggttctgctgccg<br>gacaaccactatctgtctacccagtctgcgctctctaaggacccgaacgagaaacgtgaccacatggtgctgctggagttcgtgaccgcagcgggcatcacgcacggcatggacg<br>aactgtacaaataataaCGTCTCATTAGaagcttatgcgagagtagggaactgccaggcatcaaataaaacgaaaggctcagtcgaaagactgggcctttcgttttatctgttgt<br>ttgtcggtgaacgctctcctgagtaggacaaatccgccgggagcggatttgaacgttgcgaagcaacggcccggagggtggcgggcaggacgcccgccataaactgccaggcatc<br>aaattaagcagaaggccatcctgacggatggcctttttgcgttttgcctaatgggatccaccaggcgtttaagggcaccaataactgccttaaaaaaattacgccccgccctgccac<br>tcatcgcagtactgttgtaattcattaagcattctgccgacatggaagccatcacaaacggcatgatgaacctgaatcgccagcggcatcagcaccttgtcgccttgcgtataatattt<br>gcccatggtgaaaacgggggcgaagaagttgtccatattggccacgtttaaatcaaaactggtgaaactcacccagggattggctgacacgaaaaacatattctcaataaaccctttt<br>agggaaataggccaggttttcaccgtaacacgccacatcttgcgaatatatgtgtagaaactgccggaaatcgtcgtggtattcactccagagcgatgaaaacgtttcagtttgctcat<br>ggaaaacggtgtaacaagggtgaacactatcccatatcaccagctcaccgtctttcattgccatacggaactccggatgagcattcatcaggcgggcaagaatgtgaataaaggcc<br>ggataaaacttgtgcttatttttctttacggtctttaaaaaggccgtaatatccagctgaacggtctggttataggtacattgagcaactgactgaaatgcctcaaaatgttctttacgatg<br>ccattgggatatatcaacggtggtatatccagtgatttttttctccattttagcttccttagctcctgaaaatctcgataactcaaaaaatacgcccggtagtgatcttatttcattatggtga<br>aagttggaacctcttacgtgccgatcaacgtctaattttcgccaaaagttggcccagggcttcccggtatcaacagggacaccaggatttatttattctgcgaagtgatcttccgtcac<br>aggtatttattcggcgcaaagtgcgtcgagcttatcgatgataagctgtcaaacatgagaattacaacttatatcgtatggggctgacttcaggtgctacatttgaagagataaattgca<br>ctgaaatcttgaaatattttatctgattaataagatgatcttcttgagatcgttttggtctgcgcgtaatctcttgctctgaaaacgaaaaaaccgccttgcagggcggtttttcgaaggttct<br>ctgagctaccaactctttgaaccgaggtaactggcttggaggagcgcagtcaccaaaacttgtcctttcagtttagccttaaccggcgcatgacttcaagactaactcctctaaatca<br>attaccagtggctgctgccagtggtgcttttgcatgtctttccgggttggactcaagacgatagttaccggataaggcgcagcggtcggactgaacggggggttcgtgcatacagtcca<br>gcttggagcgaactgcctacccggaactgagtgtcaggcgtggaatgagacaaacgcggccataacagcggaatgacaccggtaaaccgaaaggcaggaacaggagagcgca<br>cgagggagccgccagggggaaacgcctggtatctttatagtcctgtcgggtttcgccaccactgatttgagcgtcagatttcgtgatgcttgtcagggggggcggagcctatggaaaaa<br>cggctttgccgcggccctctcacttccctgttaagtatcttcctggcatcttccaggaaatctccgccccgttcgtaagccatttccgctcgccgcagtcgaacgaccgagcgtagcg<br>agtcagtgagcgaggaagcggaatatatcctgtatcacatattctgctgacg |

**Supplementary Table 3. Level-0 genes/parts sequences.** The sequences between BsaI sites are shown.

| Name | Sequence |
|---|---|
| **Promoter/Terminator/Insulators** | |
| $P_{BAD/araC}$ | tgaggaggattacatatgttatgacaacttgacggctacatcattcactttttcttcacaaccggcacggaactcgctcgggctggccccggtgcatttttaaatacccgcgagaaatagagttgatcgtcaaaaccaacattgcgaccgacggtggcgataggcatccgggtggtgctcaaaagcagcttcgcctggctgatacgttggtcctcgcgccagcttaagacgctaatccctaactgctggcggaaaagatgtgacagacgcgacggcgacaagcaaacatgctgtgcgacgctggcgatatcaaaattgctgtctgccaggtgatcgctgatgtactgacaagcctcgcgtacccgattatccatcggtggatggagcgactcgttaatcgcttccatgcgccgcagtaacaattgctcaagcagatttatcgccagcagctccgaatagcgcccttccccttgcccggcgttaatgatttgcccaaacaggtcgctgaaatgcggctggtgcgcttcatccgggcgaaagaaccccgtattggcaaatattgacggccagttaagccattcatgccagtaggcgcgcggacgaaagtaaacccactggtgataccattcgcgagcctccggatgacgaccgtagtgatgaatctctcctggcgggaacagcaaaatatcacccggtcggcaaacaaattctcgtccctgatttttcaccaccccctgaccgcgaatggtgagattgagaatataacctttcattcccagcggtcggtcgataaaaaaatcgagataaccgttggcctcaatcggcgttaaacccgccaccagatgggcattaaacgagtatcccggcagcaggggatcattttgcgcttcagccatacttttcatactcccgccattcagagaagaaaccaattgtccatattgcatcagacattgccgtcactgcgtcttttactggctcttctcgctaaccaaaccggtaaccccgcttattaaaagcattctgtaacaaagcgggaccaaagccatgacaaaaacgcgtaacaaaagtgtctataatcacggcagaaaagtccacattgattatttgcacggcgtcacactttgctatgccatagcatttttatccataagattagcggatcctacctgacgctttttatcgcaactctctactgtttctccatacccgtttaagcttgccaggcatcaaata |
| $P_{tac/lacI}$ | tgaggaggattacatatgtcactgcccgctttccagtcgggaaacctgtcgtgccagctgcattaatgaatcggccaacgcgcggggagaggcggtttgcgtattgggcgccagggtggtttttcttttcaccagtgaaacgggcaacagctgattgcccttcaccgcctggccctgagagagttgcagcaagcggtccacgctggtttgccccagcaggcgaaaatcctgtttgatggtggttaacggcgggatataacatgagctgtcttcggtatcgtcgtatcccactaccgagatatccgcaccaacgcgcagcccggactcggtaatggcgcgcattgcgcccagcgccatctgatcgttggcaaccagcatcgcagtgggaacgatgccctcattcagcatttgcatggtttgttgaaaaccggacatggcactccagtcgccttcccgttccgctatcggctgaatttgattgcgagtgagatatttatgccagccagccagacgcagacgcgccgagacagaacttaatgggcccgctaacagcgcgatttgctggtgacccaatgcgaccagatgctccacgcccagtcgcgtaccgtcttcatgggagaaaataatactgttgatgggtgtctggtcagagacatcaagaaataacgccggaacattagtgcaggcagcttccacagcaatggcatcctggtcatccagcggatagttaatgatcagcccactgacgcgttgcgcgagaagattgtgcaccgccgctttacaggcttcgacgccgcttcgttctaccatcgacaccaccacgctggcacccagttgatcggcgcgagatttaatcgccgcgacaatttgcgacggcgcgtgcagggccagactggaggtggcaacgccaatcagcaacgactgtttgcccgccagttgttgtgccacgcggttgggaatgtaattcagctccgccatcgccgcttccactttttcccgcgttttcgcagaaacgtggctggcctggttcaccacgcgggaaacggtctgataagagacaccggcatactctgcgacatcgtataacgttactggtttcacattcaccaccctgaattgactctcttccgggcgctatcatgccataccgcgaaaggttttgcaccattcgatgggaattcagctgttgacaattaatcatcggctcgtataatgtgtggaattgtgagcggataacaatttcacacaggaaacacaagcttgccaggcatcaaata |
| Ptet | tgaggaggattacatatgttaagacccactttcacatttaagttgtttttctaatccgcatatgatcaattcaaggccgaataagaaggctggctctgcaccttggtgatcaaataattcgatagcttgtcgtaataatggcggcatactatcagtagtaggtgtttccctttcttctttagcgacttgatgctcttgatcttccaatacgcaacctaaagtaaaatgccccacagcgctgagtgcatataatgcattctctagtgaaaaaccttgttggcataaaaaggctaattgattttcgagagtttcatactgtttttctgtaggccgtgtacctaaatgtacttttgctccatcgcgatgacttagtaaagcacatctaaaacttttagcgttattacgtaaaaaatcttgccagctttccccttctaaagggcaaaagtgagtatggtgcctatctaacatctcaatggctaaggcgtcgagcaaagcccgcttattttttacatgccaatacaatgtaggctgctctacacctagcttctgggcgagtttacgggttgttaaaccttcgattccgacctcattaagcagctctaatgcgctgttaatcactttacttttatctaatctagacatgttgacggttctccaaaacaaaaagggtacctagccactatagaccaagctgttcatttttgctttaaagccccgaaattcgattcaccttcccagctttttagttttcaatgtacgcgtccctatcagtgatagagattgacatccctatcagtgatagagatactgagcacatcagcaggacgcactgaccaagcttgccaggcatcaaata |
| $P_{J23101}$ | TTACAGCTAGCTCAGTCCTAGGTATTATGCTAGCaactttattattaaaaatta |
| $P_{J23105}$ | TTTACGGCTAGCTCAGTCCTAGGTACTATGCTAGCaactttattattaaaaatta |
| $P_{J23115}$ | TTTATAGCTAGCTCAGCCCTTGGTACAATGCTAGCaactttattattaaaaatta |
| $P_{J23117}$ | TTGACAGCTAGCTCAGTCCTAGGGATTGTGCTAGCaactttattattaaaaatta |
| $T_{L3S1P56}$ (Term) | TTTTCGAAAAAAGGCCTCCCAAATCGGGGGGCCTTTTTTATTGATAACAAAA |
| RiboJ | GCTGTCACCGGATGTGCTTTCCGGTCTGATGAGTCCGTGAGGACGAAACAGCCTCTACAAATAATTTTGTTTAA |
| **Carotenoid genes[1]** | |
| ggps* ($fds_{Y81A\_GS}$ mutant) | tgaggaggattacatATGGCGCAGCTTTCAGTTGAACAGTTTCTCAACGAGCAAAAACAGGCGGTGGAAACAGCGCTCTCCCGTTATATAGAGCGCTTAGAAGGGCCGGCGAAGCTGAAAAAGGCGATGGCGTACTCATTGGAGGCCGGCGGCAAACGAATCCGTCCGTTGCTGCTTCTGTCCACCGTTCGGGCGCTCGGCAAAGACCCGGCGGTCGGATTGCCCGTCGCCTGCGCGATTGAAATGATCCATACGATGTCTTTGATCCATGATGATTTGCCGAGCATGGACAACGATGATTTGCGGCGCGGCAAGCCGACGAACCATAAAGTGTTCGGCGAGGCGATGGCCATCTTGGCGGGGGACGGGTTGTTGACGTACGCGTTTCAATTGATCACCGAAATCGACGATGAGCGCATCCCTCCTTCCGTCCGGCTTCGGCTCATCGAACGGCTGGCGAAAGCGGCCGGTCCGGAAGGGATGGTCGCCGGTCAGGCAGCCGATATGGAAGGAGAGGGGAAAACGCTGACGCTTTCGGAGCTCGAATACATTCATCGGCATAAAACCGGGAAAATGCTGCAATACAGCGTGCACGCCGGCGCCTTGATCGGCGGCGCTGATGCCCGGCAAACGCGGGAGCTTGACGAATTCGCCGCCCATCTAGGCCTTGCCTTTCAAATTCGCGATGATATTCTCGATATTGAAGGGGCAGAAGAAAAAATCGGCAAGCCGGTCGGCAGCGACCAAAGCAACAACAAAGCGACGTATCCAGCGTTGCTGTCGCTTGCCGGCGCGAAGGAAAAGTTGGCGTTCCATATCGAGGCGGCGCAGCGCCATTTACGGAACGCTGACGTTGACGGCGCCGCGCTCGCCTATATTTGCGAACTGGTCGCCGCCCGCGACCATTAAgcttgccaggcatcaaata |
| $fds_{81A,157A\_GS}$ | tgaggaggattacatATGGCGCAGCTTTCAGTTGAACAGTTTCTCAACGAGCAAAAACAGGCGGTGGAAACAGCGCTCTCCCGTTATATAGAGCGCTTAGAAGGGCCGGCGAAGCTGAAAAAGGCGATGGCGTACTCATTGGAGGCCGGCGGCAAACGAATCCGTCCGTTGCTGCTTCTGTCCACCGTTCGGGCGCTCGGCAAAGACCCGGCGGTCGGATTGCCCGTCGCCTGCGCGATTGAAATGATCCATACGGCATCTTTGATCCATGATGATTTGCCGAGCATGGACAACGATGATTTGCGGCGCGGCAAGCCGACGAACCATAAAGTGTTCGGCGAGGCGATGGCCATCTTGGCGGGGGACGGGTTGTTGACGTACGCGTTTCAATTGATCACCGAAATCGACGATGAGCGCATCCCTCCTTCCGTCCGGCTTCGGCTCATCGAACGGCTGGCGAAAGCGGCCGGTCCGGAAGGGATGGTCGCCGGTCAGGCAGCCGATATGGAAGGAGAGGGGAAAACGCTGACGCTTTCGGAGCTCGAATACATTCATCGGCATAAAACCGGGAAAATGCTGCAATACAGCGTGCACGCCGGCGCCTTGATCGGCGGCGCTGATGCCCGGCAAACGCGGGAGCTTGACGAATTCGCCGCCCATCTAGGCCTTGCCTTTCAAATTCGCGATGATATTCTCGATATTGAAGGGGCAGAAGAAAAAATCGGCAAGCCGGTCGGCAGCGACCAAAGCAACAACAAAGCGACGTATCCAGCGTTGCTGTCGCTTGCCGGCGCGAAGGAAAAGTTGGCGTTCCATATCGAGGCGGCGCAGCGCCATTTACGGAACGCTGACGTTGACGGCGCCGCGCTCGCCTATATTTGCGAACTGGTCGCCGCCCGCGACCATTAAgcttgccaggcatcaaata |
| $crtB_{Pa}$ | tgaggaggattacatATGAATAATCCGTCGTTACTCAATCATGCGGTCGAAACGATGGCAGTTGGCTCGAAAAGTTTTGCGACAGCCTCAAAGTTATTTGATGCAAAAACCCGGCGCAGCGTACTGATGCTCTACGCCTGGTGCCGCCATTGTGACGATGTTATTGACGATCAGACGCTGGGCTTTCAGGCCCGGCAGCCTGCCTTACAAACGCCCGAACAACGTCTGATGCAACTTGAGATGAAAACGCGCCAGGCCTATGCAGGATCGCAGATGCACGAACCGGCGTTTGCGGCTTTTCAGGAAGTGGCTATGGCTCATGATATCGCCCCGGCTTACGCGTTTGATCATCTGGAAGGCTTCGCCATGGATGTACGCGAAGCGCAATACAGCCAACTGGATGATACGCTGCGCTATTGCTATCACGTTGCAGGCGTTGTCGGCTTGATGATGGCGCAAATCATGGGCGTGCGGGATAACGCCACGCTGGACCGCGCCTGTGACCTTGGGCTGGCATTTCAGTTGACCAATATTGCTCGCGATATTGTGGACGATGCGCATGCGGGCCGCTGTTATCTGCCGGCAAGCTGGCTGGAGCATGAAGGTCTGAACAAAGAGAATTATGCGGCACCTGAAAACCGTCAGGCGCTGAGCCGTATCGCCCGTCGTTTGGTGCAGGAAGCAGAACCTTACTATTTGTCTGCCACAGCCGGCCTGGCAGGGTTGCCCCTGCGTTCCGCCTGGGCAATCGCTACGGCGAAGCAGGTTTACCGGAAAATAGGTGTCAAAGTTGAACAGGCCGGTCAGCAAGC |

| | |
|---|---|
| | CTGGGATCAGCGGCAGTCAACGACCACGCCCGAAAAATTAACGCTGCTGCTGGCCGCCTCTGGTCAGGCCCTTACTTCCCGGATGCGG<br>GCTCATCCTCCCCGCCCTGCGCATCTCTGGCAGCGCCCGCTCTAGaagcttgccaggcatcaaata |
| $crtM_{Sa}$ | tgaggaggattacatATGACAATGATGAATATGAATTTTAAATATTGTCATAAAATCATGAAGAAACATTCAAAAAGCTTTTCTTACGCTTTTGACTTGTT<br>ACCAGAAGATCAAAGAAAAGCGGTTTGGGCAATTTATGCTGTGTGTCGTAAAATTGATGACAGTATAGATGTTTATGGCGATATTCAATTTTTAAA<br>TCAAATAAAAGAAGATATACAATCTATTGAAAAATACCCATATGAACATCATCACTTTCAAAGTGATCGTAGAATCATGATGGCGCTTCAGCATG<br>TTGCACAACATAAAAATATCGCCTTTCAATCTTTTTATAATCTCATTGATACTGTATATAAAGATCAACATTTTACAATGTTTGAAACGGACGCTGA<br>ATTATTCGGATATTGTTATGGTGTTGCTGGTACAGTAGGTGAAGTATTGACGCCGATTTTAAGTGATCATGAAACACATCAGACATACGATGTCG<br>CAAGAAGACTTGGTGAATCGTTGCAATTGATTAATATATTAAGAGATGTCGGTGAAGATTTTGACAATGAACGGATATATTTTAGTAAGCAACGAT<br>TAAAGCAATATGAAGTTGATATTGCTGAAGTGTACCAAAATGGTGTTAATAATCATTATATTGACTTATGGGAATATTATGCAGCTATCGCAGAAAA<br>AGATTTTCAAGATGTTATGGATCAAATCAAAGTATTTAGTATTGAAGCACAACCAATCATAGAATTAGCAGCACGTATATATATTGAAATACTGGA<br>CGAAGTGAGACAGGCTAACTATACATTACATGAACGTGTTTTTGTGGATAAGAGGAAAAAGGCAAAGTTGTTTCATGAAATAAATAGTAAATATC<br>ATAGAATATAagcttgccaggcatcaaata |
| $crtM_{AAS\_Sa}$ | tgaggaggattacatATGACAATGATGAATATGAATTTTAAATATTGTCATAAAATCATGAAGAAACATTCAAAAAGCTTTTCTTACGCTGCGGACTTGT<br>TACCAGAAGATCAAAGAAAAGCGGTTGCGGCAATTTATGCTGTGTGTCGTAAAATTGATGACAGTATAGATGTTTATGGCGATATTCAATTTTTA<br>AATCAAATAAAAGAAGATATACAATCTATTGAAAAATACCCATATGAACATCATCACTTTCAAAGTGATCGTAGAATCATGATGGCGCTTCAGCA<br>TGTTGCACAACATAAAAATATCGCCTTTCAATCTTTTTATAATCTCATTGATACTGTATATAAAGATCAACATTTTACAATGTTTGAAACGGACGCT<br>GAATTATTCGGATATTGTTATGGTGTTGCTGGTACAGTAGGTGAAGTATTGACGCCGATTTTAAGTGATCATGAAACACATCAGACATACGATGT<br>CGCAAGAAGACTTGGTGAATCGTTGCAATTGATTAATATATTAAGAGATGTCGGTGAAGATTTTGACAATGAACGGATATATTTTAGTAAGCAAC<br>GATTAAAGCAATATGAAGTTGATATTGCTGAAGTGTACCAAAATGGTGTTAATAATCATTATATTGACTTATGGGAATATTATGCAGCTATCGCAGA<br>AAAAGATTTTCAAGATGTTATGGATCAAATCAAAGTATCTAGTATTGAAGCACAACCAATCATAGAATTAGCAGCACGTATATATATTGAAATACT<br>GGACGAAGTGAGACAGGCTAACTATACATTACATGAACGTGTTTTTGTGGATAAGAGGAAAAAGGCAAAGTTGTTTCATGAAATAAATAGTAAA<br>TATCATAGAATATAagcttgccaggcatcaaata |
| $crtI_{Pa}$ | tgaggaggattacatATGAAACCAACTACGGTAATTGGTGCAGGCTTCGGTGGCCTGGCACTGGCAATTCGTCTACAAGCTGCGGGAATCCCC<br>GTCTTACTGCTTGAACAACGTGATAAACCCGGCGGTCGGGCTTATGTCTACGAGGATCAGGGGTTTACCTTTGATGCAGGCCCGACGGTTA<br>TCACCGATCCCAGTGCCATTGAAGAACTGTTTGCACTGGCAGGAAAACAGTTAAAAGAGTATGTCGAACTGCTGCCGGTTACGCCGTTTTA<br>CCGCCTGTGTTGGGAGTCAGGGAAGGTCTTTAATTACGATAACGATCAAACCCGGCTCGAAGCGCAGATTCAGCAGTTTAATCCCCGCGA<br>TGTCGAAGGTTATCGTCAGTTTCTGGACTATTCACGCGCGGTGTTTAAAGAAGGCTATCTAAAGCTCGGTACTGTCCCTTTTTTATCGTTCAGA<br>GACATGCTTCGCGCCGCACCTCAACTGGCGAAACTGCAGGCATGGAGAAGCGTTTACAGTAAGGTTGCCAGTTACATCGAAGATGAACAT<br>CTGCGCCAGGCGTTTTCTTTCCACTCGCTGTTGGTGGGCGGCAATCCCTTCGCCACCTCATCCATTTATACGTTGATACACGCGCTGGAG<br>CGTGAGTGGGGCGTCTGGTTTCCGCGTGGCGGCACCGGCGCATTAGTTCAGGGGATGATAAAGCTGTTTCAGGATCTGGGTGGCGAAGT<br>CGTGTTAAACGCCAGAGTCAGCCATATGGAAACGACAGGAAACAAGATTGAAGCCGTGCATTTAGAGGACGGTCGCAGGTTCCTGACGC<br>AAGCCGTCGCGTCAAATGCAGATGTGGTTCATACCTATCGCGACCTGTTAAGCCAGCACCCTGCCGCGGTTAAGCAGTCCAACAAACTGC<br>AGACTAAGCGCATGAGTAACTCTCTGTTTGTGCTCTATTTTGGTTTGAATCACCATCATGATCAGCTCGCGCATCACACGGTTTGTTTCGGCC<br>CGCGTTACCGCGAGCTGATTGACGAAATTTTTAATCATGATGGCCTCGCAGAGGACTTCTCACTTTATCTGCACGCGCCCTGTGTCACGGAT<br>TCGTCACTGGCGCCTGAAGGTTGCGGCAGTTACTATGTGTTGGCGCCGGTGCCACATTTAGGCACCGCGAACCTCGACTGGACGGTTGA<br>GGGGCCAAAACTACGCGACCGTATTTTTGCGTACCTTGAGCAGCATTACATGCCTGGCTTACGGAGTCAGCTGGTCACGCACCGGATGTT<br>TACGCCGTTTGATTTTCGCGACCAGCTTAATGCCTATCATGGCTCAGCCTTTTCTGTGGAGCCCGTTCTTACCCAGAGCGCCTGGTTTCGG<br>CCGCATAACCGCGATAAAACCATTACTAATCTCTACCTGGTCGGCGCAGGCACGCATCCCGGCGCAGGCATTCCTGGCGTCATCGGCTC<br>GGCAAAAGCGACAGCAGGTTTGATGCTGGAGGATCTGATATaagcttgccaggcatcaaata |
| $crtN_{Sa}$ | tgaggaggattacatATGAAGATTGCAGTAATTGGTGCAGGTGTCACAGGATTAGCAGCGGCAGCCCGTATTGCTTCTCAAGGTCATGAAGTGAC<br>GATATTTGAAAAAAATAATAATGTAGGCGGGCGTATGAATCAATTAAAGAAAGACGGCTTTACATTTGATATGGGTCCCACAATTGTCATGATGC<br>CAGATGTTTATAAAGATGTTTTTACAGCGTGTGGTAAAAATTATGAAGATTATATTGAATTGAGACAATTACGTTATATTTACGATGTGTATTTTGACC<br>ACGATGATCGTATAACGGTGCCTACAGATTTAGCTGAATTACAGCAAATGCTAGAAAGTATAGAACCTGGTTCAACGCATGGTTTTATGTCCTTT<br>TTAACGGATGTTTATAAAAAATATGAAATTGCACGTCGCTATTTCTTAGAAAGAACGTATCGCAAACCGAGTGACTTTTATAATATGACGTCACTT<br>GTGCAAGGTGCTAAGTTAAAAACGTTAAATCATGCAGATCAGCTAATTGAACATTATATTGATAACGAAAAGATACAAAAGCTTTTAGCGTTTCA<br>AACGTTATACATAGGAATTGATCCAAAACGAGGCCCGTCACTATATTCAATTATTCCTATGATTGAAATGATGTTTGGTGTGCATTTTATTAAAGG<br>CGGTATGTATGGCATGGCTCAAGGGCTAGCGCAATTAAATAAAGACTTAGGCGTTAATATTGAACTAAATGCTGAAATTGAGCAAATTATTATTG<br>ATCCTAAATTCAAACGGGCCGATGCGATAAAAGTGAATGGTGACATAAGAAAATTTGATAAAATTTTATGTACGGCTGATTTCCCTAGTGTTGC<br>GGAATCATTAATGCCAGATTTTGCACCTATTAAAAAGTATCCACCACATAAAATTGCAGACTTAGATTACTCTTGTTCAGCATTTTTAATGTATATC<br>GGTATAGATATTGATGTGACAGATCAAGTGAGACTTCATAATGTTATTTTTTCAGATGACTTTAGAGGCAATATTGAAGAAATATTTGAGGGACGTT<br>TATCATATGATCCTTCTATTTATGTGTATGTACCAGCGGTCGCTGATAAATCACTTGCGCCAGAAGGCAAAACTGGTATTTATGTGCTAATGCCG<br>ACGCCGGAACTTAAAACAGGTAGCGGAATCGATTGGTCAGATGAAGCTTTGACGCAACAAATAAAGGAAATTATTTATCGTAAATTAGCAACG<br>ATTGAAGTATTTGAAGATATAAAATCGCATATTGTTTCAGAAACAATCTTTACGCCAAATGATTTTGAGCAAACGTATCATGCGAAATTTGGTTCG<br>GCATTCGGTTTAATGCCAACTTTAGCGCAAAGTAATTATTATCGTCCACAAAATGTATCGCGAGATTATAAAGATTTATATTTTGCAGGTGCAAGT<br>ACGCATCCAGGTGCAGGCGTTCCTATTGTCTTAACGAGTGCGAAAATAACTGTAGATGAAATGATTAAAGATATTGAGCGGGGCGTATAAgctt<br>gccaggcatcaaata |
| $crtY_{Pa}$ | tgaggaggattacatATGCAACCGCATTATGATCTGATTCTCGTGGGGGCTGGACTCGCGAATGGCCTTATCGCCCTGCGTCTTCAGCAGCAGC<br>AACCTGATATGCGTATTTTGCTTATCGACGCCGCACCCCAGGCGGGCGGGAATCATACGTGGTCATTTCACCACGATGATTTGACTGAGAG<br>CCAACATCGTTGGATAGCTCCGCTGGTGGTTCATCACTGGCCCGACTATCAGGTACGCTTTCCCACACGCCGTCGTAAGCTGAACAGCG<br>GCTACTTTTGTATTACTTCTCAGCGTTTCGCTGAGGTTTTACAGCGACAGTTTGGCCCGCACTTGTGGATGGATACCGCGGTCGCAGAGGTT<br>AATGCGGAATCTGTTCGGTTGAAAAAGGGTCAGGTTATCGGTGCCCGCGCGGTGATTGACGGGCGGGGTTATGCGGCAAATTCAGCACT<br>GAGCGTGGGCTTCCAGGCGTTTATTGGCCAGGAATGGCGATTGAGCCACCCGCATGGTTTATCATCTCCCATTATCATGGATGCCACGGTC<br>GATCAGCAAAATGGTTATCGCTTCGTGTACAGCCTGCCGCTCTCGCCGACCAGATTGTTAATTGAAGACACGCACTATATTGATAATGCGAC<br>ATTAGATCCTGAATGCGCGCGGCAAAATATTTGCGACTATGCCGCGCAACAGGGTTGGCAGCTTCAGACACTGCTGCGAGAAGAACAGG<br>GCGCCTTACCCATTACTCTGTCGGGCAATGCCGACGCATTCTGGCAGCAGCGCCCCCTGGCCTGTAGTGGATTACGTGCCGGTCTGTTC<br>CATCCTACCACCGGCTATTCACTGCCGCTGGCGGTTGCCGTGGCCGACCGCCTGAGTGCACTTGATGTCTTTACGTCGGCCTCAATTCAC<br>CATGCCATTACGCATTTTGCCCGCGAGCGCTGGCAGCAGCAGGGCTTTTTCCGCATGCTGAATCGCATGCTGTTTTTAGCCGGACCCGC<br>CGATTCACGCTGGCGGGTTATGCAGCGTTTTTATGGTTTACCTGAAGATTTAATTGCCCGTTTTTATGCGGGAAAACTCACGCTGACCGATCG<br>GCTACGTATTCTGAGCGGCAAGCCGCCTGTTCCGGTATTAGCAGCATTGCAAGCCATTATGACGACTCATCGTTAAgcttgccaggcatcaaata |
| $crtZ_{Br}$ | tgaggaggattacatATGGCATGGCTGACCTGGATCGCACTGTTCCTGACCGCATTCCTGGGTATGGAGGCTTTCGCGTGGATCATGCACCGTT<br>ATGTCATGCACGGTTTCTTGTGGTCGTGGCATCGTAGCCATCACGAGCCGCACGACCACCCGCTGGAAAAGAACGACCTGTTTGCCGTTG<br>TCTTTGCCGCTCCGGCGATTGTTATGGTGGCGGTGGGTCTGCACCTGTGGCCTTGGGCCTTGCCGGTCGGTCTGGGTATTACTGCGTACG<br>GCATGGTTTACTTCTTCTTTCATGATGGCCTGGTGCATCGTCGTTTCCCGACGGGCTTTAGCGGTCGCAGCGGCTTTTGGACCCGTCGCAT<br>CCAGGCGCACCGTCTGCACCATGCAGTCCGCACGCGTGAGGGCTGCGTGTCCTTTGGCTTCTTGTGGGTTCGCAGCGCGCGTGCCCT<br>GAAAGCGGAACTGGCGCAAAAACGCGGTAGCAGCAGCTCTGGTGCATAAgcttgccaggcatcaaata |
| $crtW_{Br}$ | tgaggaggattacatATGACCGCAGCTGTCGCAGAACCTCGCATTGTACCGCGCCAAACCTGGATCGGCCTGACCCTGGCGGGTATGATTGTG<br>GCGGGCTGGGGTTCTCTGCACGTGTACGGTGTGTACTTCCACCGTTGGGGCACCAGCAGCCTGGTTATCGTCCCGGCTATCGTGGCCGT<br>TCAGACGTGGTTGTCGGTTGGCCTGTTTATTGTCGCACATGACGCCATGCACGGTTCCCTGGCCCCAGGCCGTCCGCGCCTGAACGCAG<br>CGGTGGGTCGTCTGACGCTGGGTCTGTATGCAGGCTTCCGTTTCGATCGCTTGAAAACGGCGCACCACGCGCATCATGCGGCTCCGGG<br>TACCGCAGATGACCCGGACTTTTACGCGCCTGCGCCACGCGCCTTCCTGCCGTGGTTTTTGAACTTTTTCCGTACCTATTTCGGTTGGCG<br>CGAGATGGCGGTTCTGACCGCGCTGGTCCTGATCGCGCTGTTTGGCTTGGGTGCCCGTCCGGCGAATCTGTTGACTTTTTGGGCCGCAC |

| | |
|---|---|
| | CGGCGCTGCTGAGCGCGCTGCAACTGTTCACGTTTGGCACCTGGCTGCCGCACCGTCACACGGACCAGCCGTTCGCGGATGCTCATC<br>ATGCACGCAGCAGCGGTTATGGTCCGGTTCTGAGCCTGCTGACCTGCTTTCATTTCGGTCGTCATCACGAGCACCACCTGACGCCGTGG<br>CGTCCGTGGTGGCGTTTGTGGCGTGGTGAAAGCTAAgcttgccaggcatcaaata |
| AtZEP (truncated) | tgaggaggattacatATGGCCGCAACCGCCCTGGTGGAAAAAGAAGAAAAACGTGAAGCGGTTACCGAAAAGAAAAAGAAAAGCCGTGTGCT<br>GGTTGCGGGCGGTGGCATCGGTGGCCTGGTGTTTGCCCTGGCGGCGAAAAAGAAAGGCTTTGATGTGCTGGTTTTTGAAAAAGATCTGAG<br>CGCGATTCGTGGCGAAGGCAAATATCGCGGCCCGATTCAGATTCAGAGCAATGCCCTGGCCGCACTGGAAGCCATTGATATCGAAGTGG<br>CGGAACAGGTTATGGAAGCCGGTTGCATCACCGGCGATCGCATTAATGGTCTGGTTGATGGTATTAGCGGCACCTGGTATGTGAAATTTGAT<br>ACCTTTACCCCGGCCGCAAGCCGTGGCCTGCCGGTGACCCGTGTTATCAGCCGCATGACCCTGCAGCAGATTCTGGCGCGTGCCGTG<br>GGTGAAGATGTTATCCGCAACGAAAGCAATGTGGTTGATTTTGAAGATAGCGGCGATAAAGTGACCGTGGTTCTGGAAAATGGTCAGCGCTA<br>TGAAGGCGATCTGCTGGTTGGTGCGGATGGCATTTGGAGCAAAGTGCGTAACAACCTGTTTGGTCGCAGCGAAGCCACCTATAGCGGTTA<br>TACCTGTTATACCGGCATTGCGGATTTTATCCCGGCCGATATTGAAAGCGTGGGTTATCGCGTTTTTCTGGGCCATAAACAGTATTTTGTGAGC<br>AGCGATGTTGGTGGCGGTAAAATGCAGTGGTATGCGTTTCATGAAGAACCGGCCGGCGGTGCGGATGCACCGAACGGTATGAAAAAACG<br>TCTGTTCGAAATCTTCGATGGCTGGTGCGATAATGTTCTGGATCTGCTGCATGCGACCGAAGAAGAAGCGATCCTGCGTCGCGATATTTATG<br>ATCGTAGCCCGGGCTTTACCTGGGGCAAAGGTCGCGTGACCCTGCTGGGTGATAGCATCCATGCGATGCAGCCGAACATGGGCCAGGG<br>CGGTTGTATGGCCATTGAAGATAGCTTTCAGCTGGCGCTGGAACTGGATGAAGCCTGGAAACAGAGCGTGGAAACCACCACCCCGGTTG<br>ATGTGGTTAGCAGCCTGAAACGTTATGAAGAAAGCCGTCGCCTGCGCGTTGCGATTATCCATGCGATGGCCCGTATGGCCGCAATCATGG<br>CCAGCACCTATAAAGCCTATCTGGGCGTGGGTCTGGGTCCGCTGAGCTTTCTGACCAAATTTCGTGTGCCGCATCCGGGTCGTGTGGGC<br>GGCCGTTTCTTTGTGGATATTGCGATGCCGAGCATGCTGGATTGGGTGCTGGGCGGTAATAGCGAAAAACTGCAGGGCCGTCCGCCGAG<br>CTGTCGTCTGACCGATAAAGCGGATGATCGTCTGCGCGAATGGTTTGAAGATGATGATGCCCTGGAACGTACCATCAAGGGTGAATGGTATC<br>TGATTCCGCATGGCGATGATTGCTGTGTTAGCGAAACCCTGTGCCTGACCAAAGATGAAGATCAGCCGTGTATCGTGGGTAGCGAACCGG<br>ATCAGGATTTTCCGGGCATGCGTATTGTTATCCCGAGCAGCCAGGTGAGCAAAATGCATGCGCGCGTTATTTATAAAGATGGTGCGTTTTTC<br>CTGATGGATCTGCGTAGCGAACATGGTACCTATGTGACCGATAACGAAGGCCGTCGCTATCGCGCGACCCCGAATTTTCCGGCCCGTTTT<br>CGCAGCAGCGATATTATCGAATTTGGCAGCGATAAGAAAGCGGCCTTTCGTGTGAAAGTTATTCGCAAAACCCCGAAAAGCACCCGTAAAA<br>ATGAAAGCAACAATGATAAACTGCTGCAGACCGCGTAAgcttgccaggcatcaaata |
| AtLCYb | tgaggaggattacatATGGATACCCTCCTGAAAACCCCTAACAAGCTGGACTTTTTCATCCCGCAATTCCACGGTTTCGAGCGTCTGTGCAGCAA<br>TAACCCGTACCACTCTCGTGTACGTCTGGGTGTTAAAAAGCGTGCTATTAAAATCGTTAGCTCTGTTGTGTCTGGCTCCGCAGCTCTCCTGGA<br>TCTGGTTCCGGAAACCAAGAAAGAAAACCTGGATTTCGAACTGCCGCTGTACGACACCAGCAAATCCCAGGTAGTTGATCTGGCCATCGT<br>GGGCGGTGGCCCTGCCGGCCTGGCCGTGGCACAACAGGTATCTGAAGCAGGCCTGAGCGTTTGCTCTATCGACCCATCCCCTAAACTG<br>ATCTGGCCGAATAACTACGGCGTATGGGTTGACGAATTTGAGGCAATGGATCTCCTGGACTGTCTGGACACTACGTGGAGCGGCGCGGTA<br>GTTTACGTGGATGAAGGCGTTAAAAAGGACCTGTCCCGTCCATACGGTCGCGTTAACCGCAAGCAACTGAAATCTAAAATGCTCCAGAAAT<br>GCATCACCAACGGTGTAAAGTTTCACCAGTCTAAAGTTACCAACGTCGTTCACGAAGAGGCGAACTCCACCGTCGTTTGCTCTGACGGTGT<br>CAAAATCCAGGCGTCTGTGGTTCTGGACGCGACTGGCTTCTCCCGCTGTCTGGTACAGTACGACAAACCGTACAACCCGGGCTACCAGG<br>TTGCCTACGGTATTGTTGCCGAAGTGGACGGCCACCCGTTTGATGTTGATAAAATGGTGTTCATGGACTGGCGTGACAAACACCTGGACAG<br>CTACCCGGAACTGAAAGAGCGTAACTCCAAAATCCCGACCTTCCTGTATGCGATGCCGTTCTCCTCTAACCGTATTTTTCTGGAGGAAACC<br>AGCCTGGTTGCTCGTCCTGGCCTGCGTATGGAGGACATCCAGGAACGTATGGCCGCACGCCTGAAACACCTGGGCATCAATGTGAAACG<br>CATCGAAGAGGACGAACGTTGCGTTATTCCGATGGGTGGCCCGCTGCCGGTGCTGCCGCAGCGTGTCGTGGGTATCGGTGGCACCGCA<br>GGTATGGTTCATCCGTCTACCGGCTATATGGTGGCCCGTACTCTGGCAGCGGCACCGATTGTTGCAAATGCTATTGTGCGCTACCTGGGCT<br>CCCCGTCCAGCAACAGCCTGCGTGGTGATCAGCTCTCCGCAGAAGTTTGGCGCGATCTGTGGCCTATCGAACGCCGTCGCCAGCGCG<br>AATTTTTCTGCTTTGGTATGGACATCCTCCTGAAGCTGGACCTGGATGCGACTCGTCGCTTCTTTGATGCTTTTTTCGATCTCCAGCCGCATTA<br>TTGGCATGGTTTCCTCAGCTCTCGTCTGTTTCTGCCGGAACTCCTGGTGTTCGGTCTGTCTCTGTTCTCCCATGCGAGCAACACCTCCCGTC<br>TGGAAATCATGACCAAAGGCACGGTCCCGCTGGCAAAAATGATCAATAACCTGGTCCAGGACCGCGATTAATTAagcttgccaggcatcaaata |
| AtLCYe | tgaggaggattacatATGGAATGCGTTGGTGCACGCAACTTCGCAGCGATGGCAGTTAGCACTTTCCCGTCTTGGTCTTGTCGCCGTAAATTCCC<br>GGTGGTTAAACGCTACTCCTATCGTAACATTCGCTTCGGCCTGTGCTCCGTTCGTGCATCTGGTGGCGGTTCTAGCGGTAGCGAATCTTGTG<br>TGGCGGTTCGCGAGGACTTCGCAGACGAAGAGGATTTCGTGAAAGCGGGCGGTAGCGAAATTCTGTTTGTTCAGATGCAACAGAACAAAG<br>ATATGGACGAACAGAGCAAACTGGTGGACAAACTCCCACCGATCAGCATCGGCGACGGCGCCCTGGATCACGTTGTAATCGGTTGTGGT<br>CCGGCAGGCCTGGCGCTGGCAGCCGAGAGCGCGAAGCTGGGCCTGAAGGTTGGTCTGATTGGCCCGGATCTGCCGTTCACTAATAACT<br>ATGGCGTGTGGGAGGACGAGTTTAACGACCTGGGCCTCCAGAAATGCATCGAGCACGTATGGCGTGAAACCATCGTTTATCTGGATGACG<br>ATAAACCGATTACCATTGGCCGTGCGTACGGTCGCGTGTCCCGTCGCCTCCTGCACGAGGAACTCCTGCGCCGTTGCGTTGAGAGCGGT<br>GTATCCTACCTGTCTAGCAAAGTTGATTCTATTACCGAAGCGTCCGACGGCCTGCGTCTGGTGGCCTGCGATGACAACAATGTTATCCCGTG<br>TCGCCTGGCTACGGTGGCAAGCGGCGCTGCGTCTGGCAAACTGCTCCAGTATGAAGTGGGCGGTCCGCGTGTGTGTGTGCAGACCGCA<br>TACGGTGTTGAAGTTGAGGTAGAAAATTCTCCGTACGACCCGGACCAGATGGTGTTCATGGATTATCGTGACTATACTAACGAGAAAGTGCGT<br>AGCCTCGAAGCGGAATACCCGACCTTCCTGTATGCCATGCCAATGACCAAGTCCCGCCTGTTCTTTGAGGAAACCTGTCTCGCGTCTAAA<br>GACGTGATGCCGTTTGACCTCCTGAAAACCAAACTGATGCTGCGCCTGGATACCCTGGGTATTCGTATTCTGAAAACCTACGAAGAGGAAT<br>GGTCTTATATCCCGGTCGGCGGTTCCCTGCCGAACACCGAACAGAAAAACCTGGCCTTCGGTGCAGCGGCAAGCATGGTACATCCGGC<br>GACTGGCTACTCCGTGGTTCGTTCCCTGAGCGAAGCCCCGAAATACGCTTCCGTAATCGCCGAGATCCTGCGTGAGGAAACTACCAAAC<br>AAATTAACTCCAACATTTCCCGCCAGGCTTGGGACACCCTGTGGCCTCCGGAACGCAAACGTCAGCGCGCGTTTTTCCTGTTTGGTCTGG<br>CACTGATCGTTCAGTTCGATACTGAAGGCATTCGTTCTTTTTTCCGCACTTTCTTTCGCCTGCCAAAATGGATGTGGCAAGGTTTTCTGGGTTC<br>TACCCTGACTAGCGGTGATCTGGTACTGTTCGCTCTGTACATGTTCGTTATTTCCCCAAATAACCTGCGCAAGGGTCTGATCAATCACCTGAT<br>CTCCGATCCGACCGGTGCAACCATGATCAAAACCTACCTGAAAGTGTAAttaagcttgccaggcatcaaata |
| LsLCYe (full) | tgaggaggattacatATGGAATGTTTTGGCGCGCGTAACATGACCGCCACTATGGCAGTCTTTACCTGTCCGCGTTTCACCGATTGTAACATCCG<br>TCACAAATTCTCCCTCCTGAAACAGCGCCGTTTCACCAACCTGAGCGCGTCCTCTTCCCTCCGTCAGATCAAATGCAGCGCGAAATCCGA<br>CCGCTGCGTCGTGGATAAACAGGGTATCTCTGTAGCGGACGAGGAAGATTACGTTAAAGCGGGCGGTTCCGAGCTGTTTTTCGTGCAGAT<br>GCAGCGCACTAAAAGCATGGAGTCTCAATCTAAACTGTCTGAAAAACTGGCTCAGATCCCGATTGGTAACTGTATCCTGGACCTGGTCGTTA<br>TTGGTTGTGGCCCGGCTGGCCTGGCACTGGCAGCCGAAAGCGCGAAACTGGGTCTGAACGTTGGCCTGATTGGTCCTGACCTCCCGTTT<br>ACCAATAACTATGGCGTCTGGCAGGACGAGTTCATTGGCCTGGGCCTGGAAGGTTGTATCGAACATAGCTGGAAAGACACCCTGGTTTACC<br>TGGATGACGCGGACCCGATCCGCATTGGCCGCGCTTATGGTCGTGTGCACCGCGATCTGCTCCACGAGGAACTCCTGCGTCGCTGCGT<br>GGAGTCCGGTGTGAGCTACCTCTCTTCCAAAGTCGAACGTATCACTGAAGCACCGAACGGCTACTCTCTGATCGAATGCGAAGGTAACATC<br>ACCATCCCGTGTCGTCTGGCTACCGTTGCGAGCGGTGCGGCATCTGGCAAATTCCTGGAATACGAGCTGGGTGGCCCACGTGTTTGTGT<br>GCAAACCGCCTACGGCATCGAAGTCGAAGTTGAGAATAACCCGTACGACCCTGACCTCATGGTATTTATGGATTACCGCGACTTCTCTAAA<br>CATAAACCGGAATCCCTGGAGGCGAAATACCCAACGTTCCTGTACGTGATGGCTATGTCCCCTACTAAAATCTTTTTCGAAGAGACTTGCCT<br>GGCGTCCCGTGAAGCGATGCCGTTCAACCTCCTGAAATCTAAACTGATGAGCCGTCTGAAAGCAATGGGTATCCGCATCACTCGTACCTA<br>CGAGGAAGAGTGGAGCTACATCCCGGTTGGCGGTTCCCTGCCAAACACGGAGCAGAAAAACCTGGCATTTGGTGCGGCAGCCTCTATG<br>GTGCACCCTGCGACCGGCTACTCTGTGGTTCGTAGCCTCTCTGAAGCGCCGAATTACGCCGCAGTCATTGCTAAAATTCTGCGTCAGGAT<br>CAGTCTAAGGAAATGATTTCTCTGGGTAAGTACACCAACATCTCCAAGCAGGCTTGGGAAACTCTGTGGCCGCTGGAACGTAAACGCCAG<br>CGCGCGTTCTTTCTGTTTGGCCTGTCTCACATCGTTCTCATGGACCTGGAAGGCACTCGTACCTTTTTCCGTACTTTCTTTCGTCTGCCGAAA<br>TGGATGTGGTGGGGTTTCCTGGGTTCCAGCCTGTCCTCTACCGATCTGATTATCTTCGCGCTGTACATGTTCGTTATTGCGCCGCATAGCCT<br>GCGTATGGAACTGGTGCGCCACCTCCTGTCTGATCCGACCGGCGCTACGATGGTCAAAGCGTACCTGACGATCTAATTAagcttgccaggcatc<br>aaata |
| crtU$_{Bl}$ | tgaggaggattacatATGACTCAACGCCGTCGCCCGCGTGATCGCTTCGCAGAACGTATTCAAGGCCCTCAGGGCCGTCCGCGTCTACTGCG<br>TCCGAAACGCGTAACTATCATTGGCGCAGGTATTGCTGGTCTGGCGGCAGCGGCTATTCTGGCAGAACACGGTGCAGAAGTGACCGTGAT<br>CGAAAAAAACCGATTATCTGGGCGGTCGCGTTGGTGCGTGGCCGGTTGATGACGAACGCACGATGTCTCGTGGTTTTCACGCGTTCTTTCGT |

| | |
|---|---|
| | CAGTATTACAACCTGCGTGATCTCCTGTCCCGTGCTGATCCGGAAGGTGAATGTCTGCGTCCGGTAGACGATTACCCGCTGATCCACCGC<br>CGTGGTAGCATGGACACCTTCGCTTCTATCCCGCGTACGCCACCGTTCAATCTCCTGGGCTTTGTCTGGCAGTCTCCGACGTTTCCGATCC<br>GCGGCCTGCGTGACGTGGACATCGCAGCCGCAGTCGAACTCATTGATGTGGAGTTCCCGGCCACCTACTCCTACTATGACGGCGAAAG<br>CGCAGCCGATTTTCTGGATCGTCTGCGCTTCCCGGACGAGGCGCGTCACCTGGCTCTGGAAGTGTTCGCTCGTAGCTTTTTCGCAGATCC<br>GACCGAATTTTCCGCGGGCGAACTGGTAGCTATGTTTCACACCTACTTCACCGGCAGCGCAGAAGGCCTCCTGTTCGATGTTCCGGTTGA<br>TGACTACGACACCGCTCTGTGGGCGCCGCTGGGTGGCTACCTGGAGTCTCTGGGCGTGACCATCGAAACGGGTACTACGGTTACTTCTAT<br>TGACCCAACTGAGTCCGGCTGGACTACGACCACGGGCGAAGCGAATCTCGAATCTGATGCTGTAGTGCTGGCAGTTGACCCGGCAGCG<br>GCTCGCGACCTCCTGAGCGCTTCTCACGACTCTCTGGTTGATTCTGCCCCGGCGGCTCAGCGTTGGATGGAAACCATTGGCAGCCAGAC<br>TAACGCGCCTGCTTTTGCGGTGCTGCGTCTGTGGCTCGGTACTCCGGTGGCGGACCACCGTCCTGCGTTCCTGGGCACCAGCGGTTAC<br>GACCTCCTGGACAACGTGAGCGTTCTGGAACGTTTCGAAGCAGGTGCTCGTGCATGGTCTGAATCTCATCACGGTTCTGTACTGGAACTCC<br>ACGCGTATGCACTGGAGGGTGACAGCTATGACACGGAGCGTGGTCGCGCAGATATTGTTGCACGTCTACTGTCCGACCTGCACCATGTCT<br>ATCCGGAAACGGCTGCGCTGACCATCGTAGATCAAGAACTCCTGATCGAAGCGGATTGTGGCCTGACCGACACCCGCCCGTGGGAAGA<br>TCGTCCGGAACCGTCTACTCCTATCCCGGGCCTGGTAGTTGCGGGTGACTATGTTCGCTGCAACACCCCTGTCGCGCTGATGGAACGTG<br>CTGCCACGACCGGCTACCTGGCAGCGAATCATCTCCTGTCCACCTGGCGTGTCGAGGGCACCGACCTGTGGTCCCCTCCGACCCGCG<br>GCCTCCTGCGCCGTGGTGTACTGGGCCTGATTCGCCGTCGTCGCTAATTAagcttgccaggcatcaaata |

[1]Subscript stands for; Gs: G. stearothermophillus, Pa: P. ananatis; Bl: B. linens; Mx: M. xanthus; Br: *Brevundimonas* sp. SD212

**Supplementary Table 4. Level-2 assembly.** The finalized plasmid is shown in colored background.

| Target carotenoid | Level-2 vector | pLv1.1 | pLv1.2 | pLv1.3 | pLv1.4 | pLv1.5 | pLv1.6 | End | Plasmid name |
|---|---|---|---|---|---|---|---|---|---|
| Phytoene | pACm-Lv2 | $P_{BAD/araC}$ | *ggpps** | | | | | pL1.3e-Term | pACm-Para-phy |
| 4,4'-Diapophytoene | pACm-Lv2 | pLv1.[1-2]-$P_{BAD/araC}$ | | *crtM* | | | | pL1.4e-Term | pACm-Para-C30phy |
| 4,4'-Diapo-neurosporene | pACm-Lv2 | pLv1.[1-2]-$P_{BAD/araC}$ | | *crtM* | *crtN* | | | pL1.5e-Term | pACm-Para-C30lyco |
| 4,4'-Diapolycopene | pACm-Lv2 | pLv1.[1-2]-$P_{BAD/araC}$ | | *crtM* | *crtI* | | | pL1.5e-Term | pACm-Para-C30neuro |
| C50-phytoene | pACm-Lv2 | $P_{BAD/araC}$ | *crtMaas* | *fds*$_{81,157}$ | | | | pL1.4e-Term | pACm-Para-C50phy |
| C50-b-carotene | pACm-Lv2 | $P_{BAD/araC}$ | *crtI* | *crtY* | *fds*$_{81,157}$ | *crtM*$_{AAS}$ | | pL1.6e-Term | pACm-Para-C50beta |
| δ-carotene | pACm-lyco-Lv2 | $P_{J23115}$ | *AtLCYe* | | | | | pL1.3e-Term | pACm-delta |
| ε-carotene | pACm-lyco-Lv2 | $P_{J23115}$ | *LsLCYe* | | | | | pL1.3e-Term | pACm-epsilon |
| Isorenieratene-1 | pACm-beta-Lv2 | $P_{J23115}$ | *crtU*$_{Bl}$ | | | | | pL1.3e-Term | |
| Violaxanthin-1 | pACm-zea-Lv2 | $P_{J23115}$ | *AtZEP* | | | | | pL1.3e-Term | |
| α-carotene-1 | pACm-lyco-Lv2 | $P_{J23115}$ | *AtLCYe* | *AtLCYb* | | | | pL1.4e-Term | |
| C50-zeaxanthin-1 | pACm-Lv2 | $P_{BAD/araC}$ | *crtZ*$_{Br}$ | *crtI*$_{N304P}$ | *crtY* | *fds*$_{81,157}$ | *crtM*$_{AAS}$ | pL1.7e-Term | |
| Isorenieratene-2 | pACm-beta-Lv2 | $P_{J23105}$ | *crtU*$_{Bl}$ | | | | | pL1.3e-Term | pACm-isoreni-P105 |
| Isorenieratene-3 | pACm-beta-Lv2 | $P_{tac/lacI}$ | *crtU*$_{Bl}$ | | | | | pL1.3e-Term | pACm-isoreni-Ptac |
| Violaxanthin-2 | pACm-zea-Lv2 | $P_{J23105}$ | *AtZEP* | | | | | pL1.3e-Term | |
| Violaxanthin-3 | pACm-zea-Lv2 | $P_{tac/lacI}$ | *AtZEP* | | | | | pL1.3e-Term | pACm-viola |
| α-carotene-2 | pACm-lyco-Lv2 | *AtLCYb* | $P_{J23105}$ | *AtLCYe* | | | | pL1.4e-Term | |
| α-carotene-3 | pACm-lyco-Lv2 | $P_{J23115}$ | *AtLCYe* | Term | | | | pL1.4e-AtLCYb | |
| α-carotene-4 | pACm-lyco-Lv2 | $P_{J23105}$ | *AtLCYe* | Term | | | | pL1.4e-AtLCYb | pACm-alpha |
| C50-zeaxanthin-2 | pACm-C50phy-Lv2 | $P_{BAD/araC}$ | *crtZ*$_{Br}$ | *crtI*$_{N304P}$ | *crtY* | | | pL1.5e-Term | |
| C50-zeaxanthin-3 | pACm-C50phy-Lv2 | $P_{BAD/araC}$ | RiboJ | *crtZ*$_{Br}$ | *crtI*$_{N304P}$ | *crtY* | | pL1.6e-Term | pACm-C50zea-RiboJ |
| C50-zeaxanthin-4 | pACm-C50phy-Lv2 | $P_{BAD/araC}$ | *crtI*$_{N304P}$ | *crtY* | $P_{J23105}$ | *crtZ*$_{Br}$ | | pL1.6e-Term | pACm-C50zea-P105 |
| C50-zeaxanthin-5 | pACm-C50phy-Lv2 | $P_{BAD/araC}$ | *crtI*$_{N304P}$ | *crtY* | $P_{J23101}$ | *crtZ*$_{Br}$ | | pL1.6e-Term | pACm-C50zea-P101 |

crtM$_{AAS}$: *crtM* F26A,W38A,F233S mutant
fds$_{81,157}$: *fds* Y81A,V157A mutant

### Supplementary Table 5. Primers and gene fragment used in this study.

| Primer name | Other name | Sequence |
|---|---|---|
| oMF113 | J23101 | ATAGTTTACAGCTAGCTCAGTCCTAGGTATTATGCTAGC |
| oMF114 | J23101-Rev | CGAGGCTAGCATAATACCTAGGACTGAGCTAGCTGTAAA |
| oMF115 | J23107 | ATAGTTTACGGCTAGCTCAGCCCTAGGTATTATGCTAGC |
| oMF116 | J23107-rev | CGAGGCTAGCATAATACCTAGGGCTGAGCTAGCCGTAAA |
| oMF117 | J23117 | ATAGTTGACAGCTAGCTCAGTCCTAGGGATTGTGCTAGC |
| oMF118 | J23117-rev | CGAGGCTAGCACAATCCCTAGGACTGAGCTAGCTGTCAA |
| oMF136 | J23115 | ATAGTTTATAGCTAGCTCAGCCCTTGGTACAATGCTAGC |
| oMF137 | J23115-Rev | CGAGGCTAGCATTGTACCAAGGGCTGAGCTAGCTATAAA |
| oMF206 | Lv0-vec-F | AGCTTGCCAGGCATCAAATATGTTTGAGACCAGTCAGTGGATACGCCAA |
| pMF212 | ptac-gfp-R | TATTTGATGCCTGGCAAGCTTTTATTATTT |
| oMF238 | Lv0-del-BsmBI-vecR | CATATGTAATCCTCCTCAATAGAGAGACCAAAGGGCCTCGTGATACGCCTATTTTTATA |
| oMF239 | Lv0-RBS-gfp-insF | ATTGAGGAGGATTACATATGAATTCAGCTGTTGACAATTAATCATCGG |
| oMF273 | pZERO-vecF | CTGACGTCTAAGAAACCATTATTATCATGA |
| oMF274 | pZERO-vecR | TTCAAATATGTATCCGCTCATGAGACAATA |
| oMF275 | Term-F | GCGGATACATATTTGAAATGCGAGAGTAGGGAACTGC |
| oMF276 | Term-R | GTTTCTTAGACGTCAGCGGCGGATTTGTCCTACTCA |
| oMF367 | BsmBI-F | gctggaggatctgatatgaaGGGCCCATCTTGAGACGtgacaattaatcatcggctcgtataatgt |
| oMF368 | BsmBI-R | tccctactctcgcataagcttCTAATGAGACGTTATTATTTGTACAGTTCG |
| gblock #103986008 | lacZ-casette | CAATTTCACACAGGAAACACTCGAGTACGTCTCATGCCCTATAGAGACCTTACAGCTAGCTCAGTCCTAGGTATTATGCTAGCGCTATGACCATGGTACAAAGAGGAGAAAGGACATGGGTGACCCGAAACTGCTGCTGGAAGTTCCGCACGCTATCTCTTCTTCTCCGGGTAACTCTCTGGCTGTTGTTCTGCAACGTCGTGACTGGGAAAACCCGGGTGTTACCCAGCTGAACCGTCTGGCTGCTCACCCGCCGTTCGCTTCTTGGCGTAACTCTGAAGAAGCTCGTACCGACCGTCCGTCCCAGCAGCTGCGTTCTCTGATCCGTCTGCTGACCAAACCGGAACGTAAACTGTCTTGGCTGCTGCCGCCGCTGTCTAACAACTAACGCAAAAAAGGTCTCATGTTCCGGAGAGACGCAAGCTTGCCAGGCATCAAATAAAAC |

**Supplementary Note 1. Strategies to construct "basic" carotenoid operons.**

(1) Operon expression from a single promoter. Using multiple terminators and promoters can make it difficult to introduce other plasmids and may destabilize plasmid retention due to the use of the same genetic parts (DNA sequence).

(2) The relative expression levels of the genes is known to be somewhat, if not perfectly,[1-3] controlled by the location of the genes within the operon,[4] where the front gene being highest and the last being lowest expression. Thus, genes located in front of the operon (closer to the promoter) can be assumed to have higher expression levels compared to the downstream genes.

(3) To make the production levels roughly the same for all constructs, GGPS* and CrtB (phytoene synthesis, which determines the flux towards carotenoid synthesis) are placed at the beginning of the operon. The order of the genes placed after them is arranged downstream of the pathway. For example, in the case of Zeaxanthin, it is arranged as ggpps-crtB followed by crtZ-crtY-crtI.

(4) For the GGPP synthase: instead of using the Pantoea-derived CrtE which is commonly used as GGPP synthase, use the *G. stearothermophillus* $FDS_{Y81M}$ mutant (renamed GGPS* in this paper). While CrtE uses FPP and IPP as substrates, GGPS* can use FPP, IPP and also DMAPP as substrates. This is considered more optimal for carotenoid production as it is not dependent on the host's FPP levels.


1. Burkhardt, D. H. *et al.* Operon mRNAs are organized into ORF-centric structures that predict translation efficiency. *eLife* **6**, e22037 (2017).
2. Jeschek, M., Gerngross, D. & Panke, S. Combinatorial pathway optimization for streamlined metabolic engineering. *Current Opinion in Biotechnology* **47**, 142–151 (2017).
3. Gerngross, D., Beerenwinkel, N. & Panke, S. Systematic investigation of synthetic operon designs enables prediction and control of expression levels of multiple proteins. 2022.06.10.495604 Preprint at https://doi.org/10.1101/2022.06.10.495604 (2022).
4. Lim, H. N., Lee, Y. & Hussein, R. Fundamental relationship between operon organization and gene expression. *Proc Natl Acad Sci U S A* **108**, 10626–10631 (2011).
